# SUPREME: A cancer subtype prediction methodology integrating multiomics data using Graph Convolutional Neural Network

**DOI:** 10.1101/2022.08.03.502682

**Authors:** Ziynet Nesibe Kesimoglu, Serdar Bozdag

## Abstract

To pave the road towards precision medicine in cancer, patients with highly similar biology ought to be grouped into the same cancer subtypes. Utilizing high-dimensional multiomics datasets, several integrative computational approaches have been developed to uncover cancer subtypes. Recently, Graph Neural Networks (GNNs) was discovered to learn node embeddings while utilizing node features and node associations at the same time on graph-structured data. Although there are some commonly used architectures such as Graph Convolutional Network (GCN) for cancer subtype prediction, the existing prediction tools have some limitations in leveraging those architectures with multiomics integration on multiple networks. Addressing them, we developed SUPREME (a <u>su</u>btype <u>pre</u>diction <u>me</u>thodology) by comprehensively analyzing multiomics data and associations between patients with graph convolutions on multiple patient similarity networks. Unlike the existing tools, SUPREME generates patient embeddings from patient similarity networks, on which it utilizes all the multiomics features. In addition, SUPREME integrates all the possible combinations of embeddings with the raw multiomics features to capture the complementary signals. Extensive evaluation of all combinations makes SUPREME interpretable in terms of utilized networks and features. On three different datasets from The Cancer Genome Atlas (TCGA), Molecular Taxonomy of Breast Cancer International Consortium (METABRIC), and both combined, our method significantly outperformed other integrative cancer (sub)type prediction tools and baseline methods, with overall consistent results. SUPREME-inferred subtypes had significant survival differences, mostly having more significance than ground truth (PAM50) labels, and outperformed nine cancer subtype differentiating tools and baseline methods. These results suggest that with proper utilization of multiple datatypes and patient associations, SUPREME could demystify the undiscovered characteristics in cancer subtypes that cause significant survival differences and could improve the ground truth label, which depends mainly on a single datatype. Source code for our tool is publicly available at https://github.com/bozdaglab/SUPREME.

## 1 Introduction

Cancer is one of the deadliest diseases for which cancer-causing agents such as oncogenes, mutations, and gene regulatory associations have not been fully demystified. Cancer patients show different characteristics in terms of the progression of disease and response to treatment [1]. Various biological datasets from cancer tissues have been generated to better characterize cancer biology. For instance, The Cancer Genome Atlas (TCGA) project generated over 2.5 petabytes of multiple omics (multiomics) data for thousands of patients from 33 different cancer types (available at https://www.cancer.gov/tcga). Specifically for breast cancer, the Molecular Taxonomy of Breast Cancer International Consortium (METABRIC) has generated four types of multiomics data for thousands of breast tumor samples [2]. Utilizing the high-dimensional biological datasets in public databases, computational approaches have been developed to discover subtypes of various cancers [3, 4, 5]. Several of the cancer subtype prediction studies rely only on one type of biological datatype [4, 6, 7]. However, each of these datatypes captures a different part of the underlying biology, thus developing integrative computational methods has been an important research area in bioinformatics.

Breast cancer is currently the most commonly-diagnosed cancer worldwide [8]. Therapeutic groups in breast cancer (i.e., estrogen receptor-positive, progesterone receptor-positive, human epidermal growth factor receptor 2 (HER2) amplified group, and triple-negative breast cancer) mainly depend on three receptors. Even though these receptors are very impactful in determining the breast cancer subtypes, they are not solely sufficient to classify a patient. Some other studies showed the genomic and clinical features such as race, age, and some mutations are also important in breast cancer subtyping [9, 10].

Genomic datatypes are found informative for differentiating subgroups in breast cancer. In 2009, Parker et al. [11] found a clear difference in the expression of 50 genes for breast cancer and introduced breast cancer molecular subtypes, called *PAM50 subtypes*. In 2012, the TCGA group published a study analyzing breast cancer subgroups and their associations with single datatypes, obtaining subtype-specific patterns in each datatype [12] and supporting the importance of gene expression like the PAM50 model [11]. Even though there are important signals from both clinical and genomic features to determine the subtype of a patient, relying on a single data modality is not sufficient to differentiate subtypes clearly. As we get more samples and datatypes to analyze, it is important to integrate all the available datatypes properly with advanced approaches to understand differences in the characteristics of cancer patients.

Recently several groups have developed unsupervised computational tools to integrate multiple datatypes to discover cancer subtypes. For instance, iClusterPlus [13] uses a joint latent variable model concatenating multiple datatypes with dimension reduction to cluster cancer patients. Similarity Network Fusion (SNF) [14] builds a patient similarity network based on each datatype, obtains a fused patient network by applying a nonlinear fusion step, and performs the clustering on that final network. PINSPlus [15] assumes that samples that are truly in the same subtype are clustered together despite small changes in the data. PINSPlus discovers the subtypes if the samples are highly connected for different datatypes applying data perturbation. The authors demonstrated that PINSPlus had robust results with significant survival differences across different cancer types. Those studies focus on unsupervised multiomics data integration without the utilization of found subtype labels such as PAM50 subtype labels. Furthermore, these tools utilize patient similarity networks or features, but not both simultaneously, while there are recent improvements in graph representation learning allowing to utilize both at the same time [16, 17, 18].

Graphs (networks) are suitable data structures to store multiomics datasets. Deep learning-based architectures have been used extensively for grid-like data (e.g. image), however, these methods are not directly applicable to graph data. Graphs are unstructured as each node has a varying number of neighbors and there is no fixed ordering of nodes. To train models on graph data, some shallow learning methods emerged by encoding every node into a fixed low-dimensional vector, called *embedding*, representing the position and the local relationships in the graph [19, 20, 21]. However, these embedding methods are not scalable for large graphs and cannot utilize the node features that we have plenty of, thus, current methods are being replaced with more advanced deep learning-based methods such as Graph Neural Network (GNN) [16, 17]. GNNs have recently been applied to biological problems such as cancer type/subtype prediction and drug response prediction [18, 22, 23, 24, 25]. The main difference in GNN-based architectures is how the features are aggregated from the local structure. Graph Convolutional Networks (GCN) is one of the most popular GNNs that uses a modified aggregation involving self-nodes with normalization across neighbors [18]. Even though there are some studies applying convolution to graph-structured data for cancer subtyping, these models are mostly applicable to a single network or had some limitations for integrative approaches.

In [26], cancer type prediction of patients from 33 cancer and non-cancer types (i.e., all normal samples from all 33 available cancer types) was performed using GCNs. The input network was based on gene coexpression or protein-protein interaction, but the convolution was done on the gene expression dataset only, thus, missing the information of multiple data modalities. Multiomics GCN (MOGONET) is a supervised multiomics integration framework using GCNs with a patient similarity network for mRNA expression, DNA methylation, and microRNA expression separately [27]. MOGONET gets the label independently from three different models, then uses them to get the final prediction. However, it does not consider multiple features for networks. We call this kind of embedding *datatype-specific patient embedding* where the methodology generates datatype-specific networks with datatype-specific node features and considers only the prediction labels from separate GCN models. However, these embeddings could be improved by utilizing all the multiomics patient features on each local network structure, making the embedding *network-specific patient embedding*. Moreover, it is possible to utilize GCN not only to get the prediction label but also to obtain the embeddings and integrate them. Going further, we can also integrate the patient features themselves (called *raw features*) with embeddings to capture any diluted features, which themselves play an important role in the task. To utilize more from available knowledge, it is important to properly integrate multiple network representations and multiomics features simultaneously.

To address the aforementioned limitations, we developed a computational tool named SUPREME, a <u>su</u>btype <u>pre</u>diction <u>me</u>thodology, integrating multiple types of biological datasets using GCNs. SUPREME generates patient similarity networks using all available datatypes and convolves the multiomics features on each network assuming that patients with a similar local neighborhood are likely to belong to the same subtype, where the similarity is based on a specific datatype’s features. SUPREME encodes the relations on patient similarity network from each biological datatype to consider intra-datatype relation and obtains network-specific patient embeddings incorporating patient multiomics features on each network to consider inter-datatype relation. Then SUPREME integrates these embeddings providing extensive evaluations of all combinations of patient embeddings. For each combination, SUPREME integrates the selected patient embeddings with raw multiomics features to utilize all the knowledge at the same time. SUPREME utilizes all available datatypes from public datasets and can interpret each datatype’s effectiveness in terms of features and networks. Being model-agnostic, SUPREME could be easily adapted to any model, any prediction task handling any number of datatypes, and could be easily modified by changing the Machine Learning (ML) method, network generation strategy, and feature extraction approach.

We applied SUPREME to the breast cancer datasets from TCGA and METABRIC databases separately and together. Our results on the cancer subtype prediction task showed that SUPREME outperformed other integrative supervised cancer (sub)type prediction tools and baseline methods. SUPREME had improved performance showing the importance of GCN-based approaches, network-specific patient embeddings, and raw feature integration. SUPREME was robust showing high and consistent prediction performance. We observed that the gene expression (EXP)-based features were the most significant features, as expected for breast cancer. Importantly, SUPREME-inferred cancer subtypes had consistently significant survival differences in cancer subtypes and were mostly more significant than the survival differences between the ground truth subtypes, which were based on gene expression datatype. These results suggest that SUPREME can differentiate the characteristics of cancer subtypes properly utilizing the multiple network relations and multiple datatypes.

## 2 Results

### 2.1 SUPREME Framework & Evaluation

We introduced a novel node classification framework, called SUPREME (a <u>su</u>btype <u>pre</u>diction <u>me</u>thodology), that utilizes graph convolutions on multiple datatype-specific networks that are annotated with multiomics datatypes as node features (Figure 1). This framework is model-agnostic and could be applied to any classification problem with properly processed datatypes and networks. In this work, SUPREME was applied specifically to the breast cancer subtype prediction problem by applying convolution on patient similarity networks constructed based on multiple biological datatypes from breast tumor samples.

**Figure 1.**
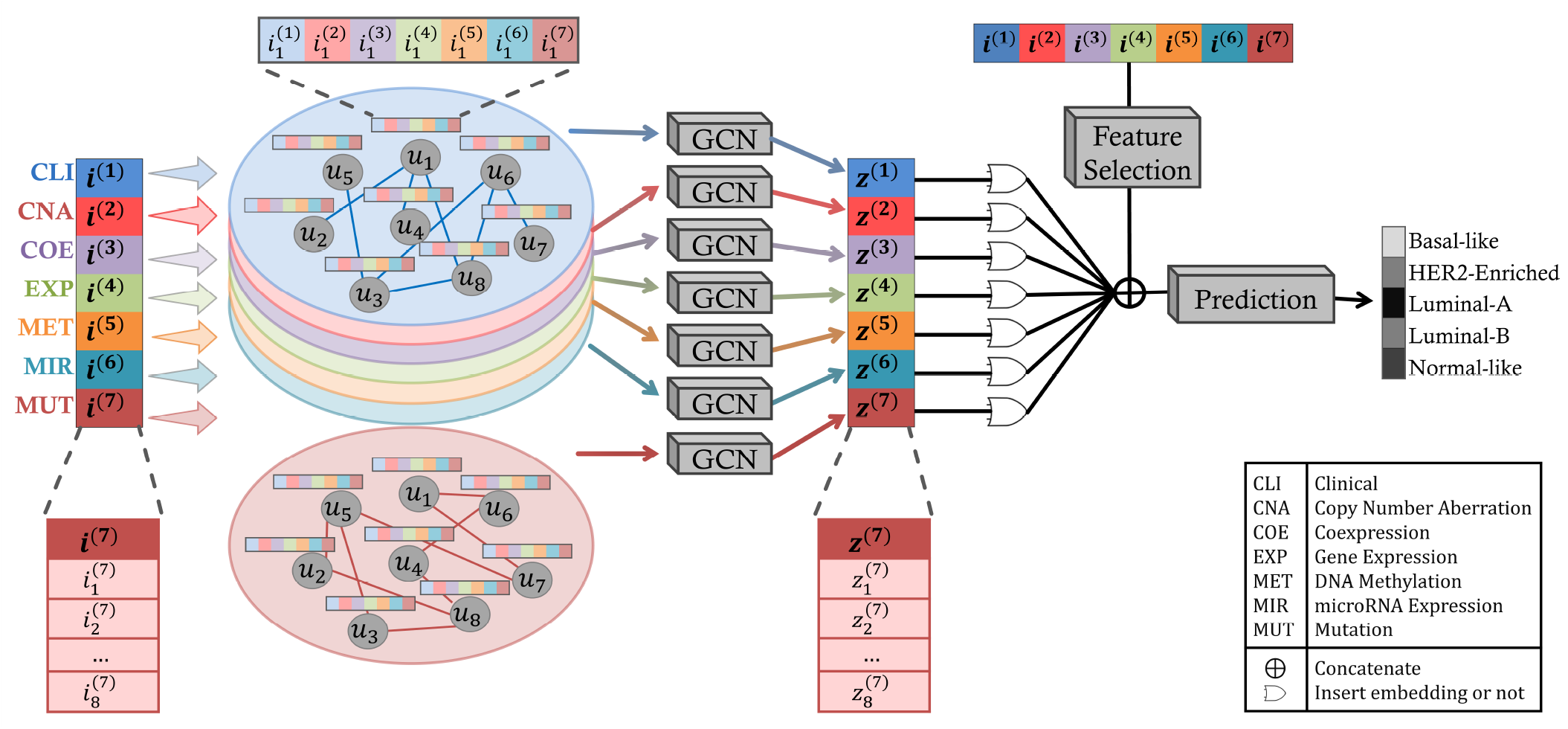
The SUPREME Pipeline. SUPREME generates patient similarity networks using all available datatypes where nodes are annotated with multiomics features. Utilizing graph convolutions on each patient similarity network, patient embeddings are generated. To provide extensive evaluations of subtype prediction, a machine learning model is trained for each combination of patient embeddings and raw multiomics features. [*u*_*k*_ is *k*^*th*^ patient, *i*^(*j*)^ is a raw feature matrix for the *j*^*th*^ datatype where each row is 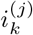 corresponding the feature vector of the *k*^*th*^ patient for *j*^*th*^ datatype. Similarly, *z*^(*j*)^is a node embedding matrix for *j*^*th*^ datatype-specific network where each row is 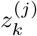 corresponding the embedding of the*k*^*th*^ patient.]

We applied SUPREME on three datasets for the breast cancer subtype prediction task. We collected the data and generated seven datatypes (i.e., clinical, copy number aberration, coexpression, gene expression, DNA methylation, microRNA expression, and mutation) across 1022 breast tumor samples from TCGA [12], five datatypes (i.e., clinical, copy number aberration, coexpression, gene expression, and mutation) across 1699 breast tumor samples from METABRIC [2] and three datatypes (clinical, gene expression, and mutation) across a total of 2721 breast tumor samples from the combined datasets of TCGA and METABRIC. As ground truth for the prediction task, we obtained the PAM50 subtype labels, namely Basal-like, HER2-Enriched, Luminal-A, Luminal-B, and Normal-like [11]. We preprocessed data and extracted features from each datatype separately (i.e., raw multiomics features). We used these features to generate patient similarity networks, used them as node features in the patient similarity networks and also integrated them with patient embeddings during the subtype prediction task. Table 1 shows the number of samples, subtypes, and features for each datatype in each dataset.

**Table 1.** Number of features and samples for each dataset. Subtypes are abbreviated as BL: Basal-like, HER2: HER2-Enriched, LA: Luminal-A, LB: Luminal-B, NL: Normal-like. See Figure 1 for the abbreviations of datatypes.

| Dataset | Number of raw features | Number of samples |
| --- | --- | --- |
| TCGA | 29 CLI, 500 CNA, 16 COE, 1000 EXP, 1000 MET, 343 MIR, 200 MUT | 1022 samples: 172 BL (17%), 78 HER2 (8%), 538 LA (53%), 195 LB (19%), 39 NL (4%) |
| METABRIC | 29 CLI, 500 CNA, 32 COE, 1000 EXP, 200 MUT | 1699 samples: 199 BL (12%), 220 HER2 (13%), 679 LA (40%), 461 LB (27%), 140 NL (8%) |
| Combined | 29 CLI, 1000 EXP, 200 MUT | 2721 samples: 371 BL (14%), 298 HER2 (11%), 1217 LA (45%), 656 LB (24%), 179 NL (7%) |

We computed patient embeddings training a GCN on each patient similarity network separately. Different than the studies utilizing a single datatype on GCN, we trained each patient similarity network separately but included all raw features of the patients as node features in each network to grasp the complementary signals from multiple datatypes. Thus, we generated a network-specific patient embedding per datatype (seven for TCGA data, five for METABRIC, and three for the combined data) by exploiting the information from multiomics features and network topology simultaneously using GCN.

To predict cancer subtypes, we trained ML models integrating patient embeddings with the raw multiomics features to capture their complementary signals. To show the overall performance of the methods regardless of the selected networks, we trained a prediction model for each combination of patient embeddings, having a total of 2^𝕕^ − 1 different models for each SUPREME evaluation, where 𝕕 is the number of the datatypes. In that way, we had multiple representations and were able to measure the effectiveness of patient embeddings and features from each datatype. Specifically, we had 127 different prediction models for TCGA data, 31 for METABRIC data, and seven for the combined data. To see the impact of raw feature integration to the embeddings, we also trained models without integrating raw features with patient embeddings, called *SUPREME-*.

SUPREME builds two types of models: a GCN model to learn patient embeddings and a machine learning model for the final subtype prediction. We used 60% and 20% of the samples for training and validation of the models, respectively. We hold out the remaining 20% as the test set and never used them except for the final evaluation of the tool. This splitting was done as stratified, i.e., keeping the same ratio of the subtype labels in the original data for each split to deal with imbalanced class sizes. To tune the hyperparameters, for each hyperparameter combination, SUPREME trained a model using the training data, measured the macro-averaged F1 (macro F1) score on the validation data 10 times (See Supplemental File 1) and used the hyperparameter combination giving the best median macro F1 score to generate the final model (See Supplemental File 2 and Supplemental Figure 1 for the selected hyperparameters and the training/validation details).

To evaluate the performance of SUPREME and baseline methods, we measured the macro F1 score, accuracy, and weighted-average F1 (weighted F1) score. We used the violin plot to visualize the performance results across all embedding combinations. For the prediction task, we tested four ML algorithms, namely Multi-layer Perceptron (MLP), Random Forest, Support Vector Machine, and XGBoost (Supplemental Figure 2), and decided to use MLP due to its consistently high performance (Supplemental Table 1 and discussion for details).

### 2.2 SUPREME outperformed the cancer subtype prediction tools and baseline methods

We compared the performance of SUPREME on three different datasets with seven other cancer (sub)type prediction tools and baseline methods, namely Deep cancer subtype classification (DeepCC) [28], GCN-based classification (GCNC) [26], MOGONET [27], MLP, Random Forest, Support Vector Machine, and XGBoost. MOGONET utilizes GCN on multiomics data utilizing datatype-specific embedding predictions. GCNC leverages GCN with gene expression features on protein-protein interaction (PPI)- or coexpression-based gene network, while DeepCC utilizes only gene expression datatype with pathway activity transformation through an MLP model. Therefore, we had only two classification models for GCNC: *GCNC* _*PPI*_ with the PPI network and *GCNC* _*COE*_ with the coexpression network, and one model for DeepCC. Other than DeepCC and GCNC, we ran SUPREME, SUPREME-, and the other tools for all the combinations of available datatypes.

SUPREME and SUPREME-outperformed all other multi-omics integration methods for three datasets in terms of macro F1, accuracy, and weighted F1 (Figure 2 and Supplemental Figures 3 and 4). SUPREME significantly outperformed MLP, which utilizes raw features only in all datasets, showing the importance of the GCN utilization. In METABRIC and the combined data, we observed that SUPREME significantly outperformed SUPREME-. In TCGA data, 76% of the SUPREME models had higher macro F1 than their corresponding SUPREME- models had, showing the importance of raw feature integration.

**Figure 2.**
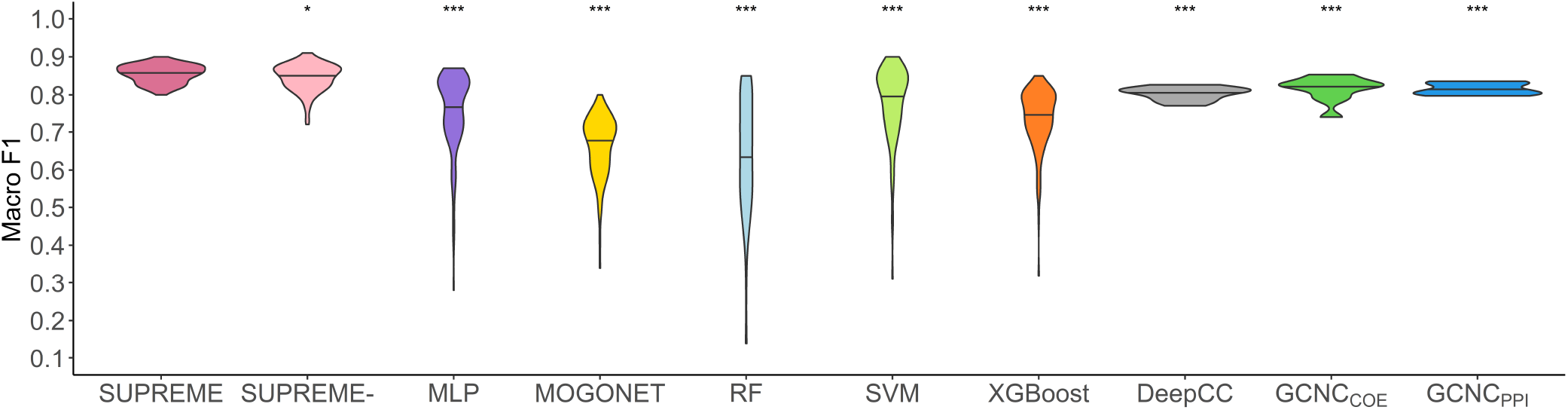
Classification results. Violin plot of macro F1 scores obtained from 127 different models including all different combinations of datatypes as compared to the cancer subtype prediction tools and baseline supervised methods on TCGA data. DeepCC and GCNC violin plots show the distribution of macro F1 scores of ten runs of a single model as they can only utilize gene expression datatype. The significance level was measured with respect to SUPREME (Wilcoxon rank-sum test p-value to compare the distribution of violin plots representing the significance *<* 0.001 by ***, else if *<* 0.01 by **, and else if *<* 0.05 by *). [MLP: Multi-layer Perceptron, RF: Random Forest, SUPREME-: SUPREME without raw feature integration, SVM: Support Vector Machine]

We ran the tools that utilize only gene expression datatype and evaluated their performance (Supplemental Table 2). For TCGA data, SUPREME achieved significantly higher performance than DeepCC and GCNC models (Figure 2), while performance on the combined data was comparable (Figure 2) and Supplemental Figure 3).

**Table 2.** Single model results on TCGA data. Macro F1 scores for each model with a single dataype. See Figure 1 for the abbreviations of the datatypes. [MLP: Multi-layer Perceptron.]

| Method | CLI | CNA | COE | EXP | MET | MIR | MUT |
| --- | --- | --- | --- | --- | --- | --- | --- |
| SUPREME | 0.68±0.04 | **0.80±0.03** | 0.76±0.04 | **0.84±0.02** | **0.79±0.03** | 0.73±0.02 | **0.75±0.03** |
| SUPREME- | **0.72±0.02** | 0.77±0.02 | **0.77±0.04** | 0.77±0.05 | **0.79±0.04** | 0.70±0.02 | 0.74±0.04 |
| MLP | 0.46±0.07 | 0.53±0.04 | 0.59±0.02 | 0.82±0.03 | 0.69±0.04 | **0.74±0.04** | 0.28±0.06 |
| MOGONET | 0.41±0.01 | 0.52±0.01 | 0.57±0.01 | 0.75±0.01 | 0.61±0.03 | 0.71±0.03 | 0.34±0.01 |

In addition, we checked the subtype-specific F1 scores, and had consistent and higher performance across all subtypes, mostly having significant differences (Supplemental Figure 5). Specifically, on TCGA data, we had significantly better performance than all other tools for all subtypes in terms of subtype-specific F1 scores. Particularly, SUPREME had a significantly higher subtype-specific F1 score than all other tools on the Normal-like subtype for all three datasets. Considering that the Normal-like subtype had the smallest sample size in all three datasets (4% of the samples from TCGA, 8% from METABRIC, and 7% from the combined data), achieving this performance increase indicates SUPREME’s robustness even for minority classes.

### 2.3 SUPREME had consistently high performance even with single models

To see SUPREME’s performance with a single datatype, we investigated models generated with only one datatype, called *single model*. We compared SUPREME with an MLP-based model trained using a single datatype to show the impact of our GCN-based approach. To show the impact of different approaches with one datatype, we compared our single models against MOGONET, and our EXP-based model with DeepCC and GCNC models. We conducted these experiments for all three datasets.

Based on the single model results, SUPREME outperformed MOGONET for all single models from all three datasets (Table 2, and Supplemental Tables 3, 4 and 5). Also, SUPREME outperformed MLP (six out of seven models for TCGA data, three out of five for METABRIC data, and two out of three models for the combined data), or had comparable performance, while MLP had extremely poor performance on some datatypes, showing the importance of GCN-based approach.

**Table 3.**
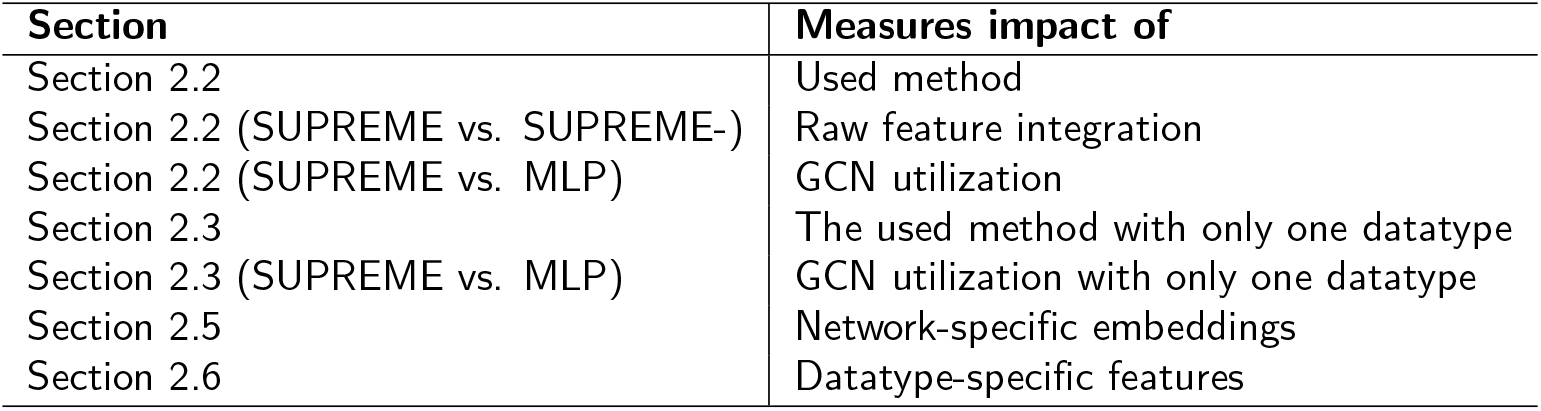
Ablation Studies.

There was no clear winner for the comparison of the SUPREME EXP-based model with DeepCC and GCNC models. In terms of macro F1 score, SUPREME outperformed both methods on TCGA data and GCNC (1 draw, 1 win) on the combined data. (Table 2, and Supplemental Tables 3, 4 and 5). This could be because DeepCC and GCNC utilize pathway activation, PPI network, or coexpression network in addition to gene expression datatype. Nonetheless, by utilizing more datatypes SUPREME outperformed or was on par with both tools for all datasets (Supplemental Figures 3 and 4).

EXP-based models had the highest macro F1 score for all three datasets for all methods (Table 2, and Supplemental Tables 3, 4 and 5). The only exception is that SUPREME-MET-based model had slightly higher performance than SUPREME- EXP-based model on TCGA data. High performance of EXP-based models is not surprising as the breast cancer subtype labels are based on gene expression data. We observed that SUPREME usually outperformed SUPREME- on single models, which indicates that utilizing raw features usually improve the model performance. On the other hand, there were few cases where adding raw features dropped the performance (e.g., CLI-based models on TCGA data). By examining SUPREME- and MLP model performances, we compared the predictive power of patient embeddings with raw features. We observed that patient embedding features were more useful than raw features with few exceptions, such as microRNA expression- (MIR) and EXP-based models on TCGA data, copy number aberration (CNA)-based on METABRIC data, and CLI-based model on the combined data. Specifically on TCGA, we see that CLI-based embedding was more informative than CLI-based features. For CNA- and mutation (MUT)-based models, embeddings were more useful than raw features, but we observed that integrating raw features to embeddings further improved the performance. Similarly, although for EXP-based model on TCGA data, embeddings were less informative than raw features, integrating them improved the performance.

### 2.4 SUPREME had significant survival differences between predicted subtypes consistently

To measure the ability of the supervised methods to differentiate samples based on survival, we predicted the subtype labels for each data modality combination and performed the survival analysis. In addition to the supervised methods, we also included the state-of-the-art unsupervised tools that are specifically applied to cancer subtyping (i.e., iClusterPlus [13], Similarity Network Fusion (SNF) [14], and PINSPlus [15]) and an algorithmically-relevant clustering method (i.e., Affinity Propagation (AP) clustering). AP clustering is relevant because it uses a message-passing strategy to find the cluster representatives and the best representative for each node. We obtained five clusters from the unsupervised methods to match the number of PAM50 subtypes and checked the survival differences for these obtained clusters. This analysis was only applied for the results on TCGA data where patient survival data were available. To check the statistical significance of survival difference between subtypes, we applied the log-rank test to compute p-values.

The results showed that SUPREME’s predicted subtypes consistently had significant differences in survival rates and significantly outperformed all other nine methods in terms of the p-value (Figure 3). SUPREME had 0.0035 as the lowest p-value (when integrating CNA-, COE-, MET-, and MUT-based patient embeddings) and 0.0131 as the median p-value (Supplemental Figure 6A for the Kaplan-Meier plot). Similarly for SUPREME-, we had 0.0018 as the lowest p-value (when integrating CNA- and COE-based patient embeddings), and 0.0147 as the median p-value. Interestingly, SUPREME had a more significant survival difference than survival difference between ground truth (i.e., PAM50) labels (Supplemental Figure 6B for the Kaplan-Meier plot for PAM50 subtypes).

**Figure 3.**
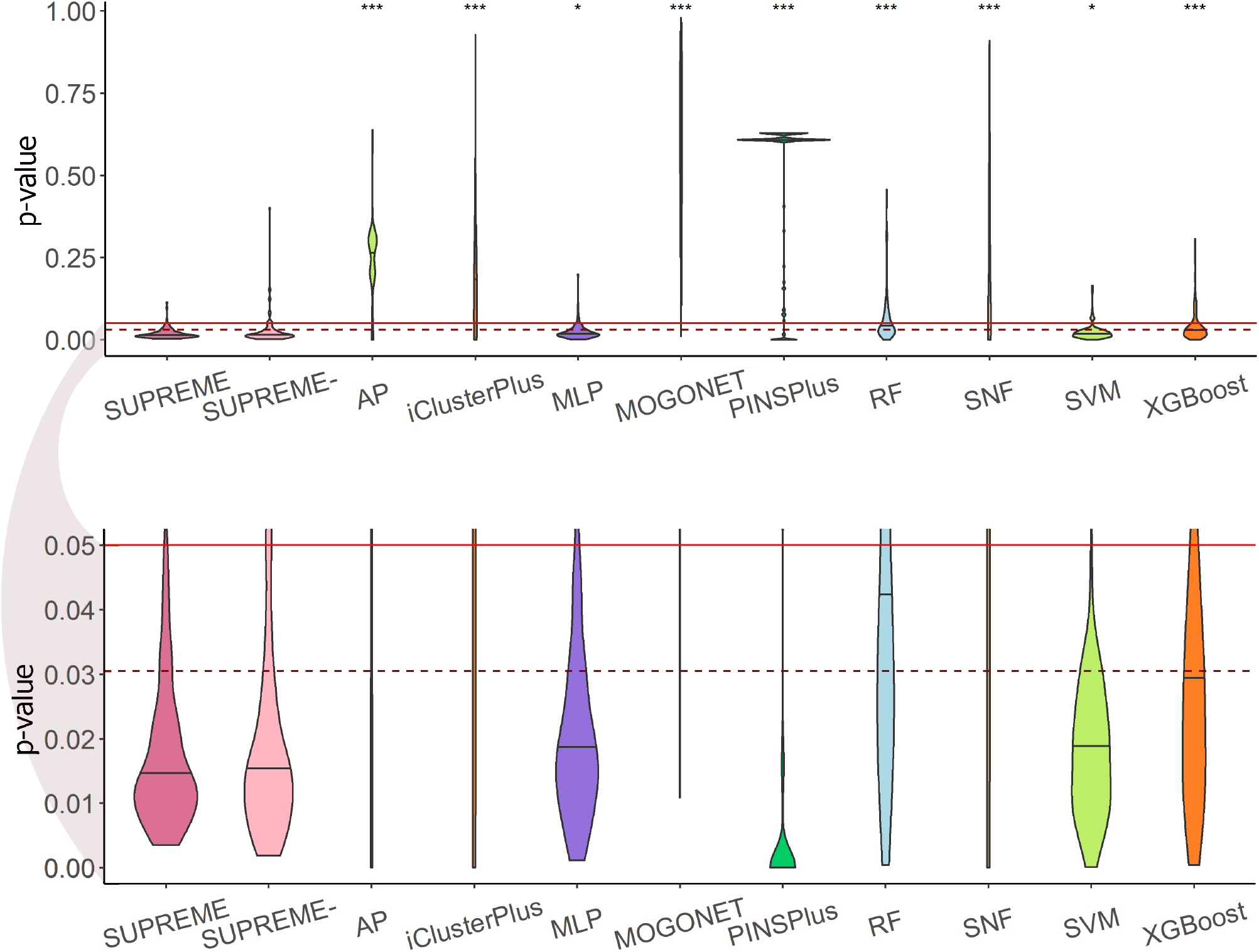
Survival Analysis Results. Violin plot of the log-rank p-value obtained from survival analysis for the SUPREME models as compared to the cancer subtype prediction/clustering tools and baseline methods. Significance level was measured with respect to SUPREME (Wilcoxon rank-sum test p-value to compare the distribution of violin plots representing the significance *<* 0.001 by ***, else if *<* 0.01 by **, and else if *<* 0.05 by *). The continuous line shows the significance level of 0.05 and the dashed line shows the ground truth’s significance level. The below figure focuses on the significant survival p-values (*<* 0.05). [AP: Affinity Propagation, MLP: Multi-layer Perceptron, RF: Random Forest, SNF: Similarity Network Fusion, SUPREME-: SUPREME without raw feature integration, SVM: Support Vector Machine]

Specifically, 106 out of 127 SUPREME models had a lower p-value than the p-value for ground truth. For 57% of those models, we had CNA-based embedding selected. It is followed by 52% from COE-, CLI-, and MET-based embeddings. This might suggest that those embeddings could contribute more to differentiating survival differences between subtypes.

AP, iClusterPlus, MOGONET, and SNF methods had a wide range of p-values, while SUPREME, MLP, SVM, and XGBoost had mostly significant p-values (*≤* 0.05) with a median lower than the significance level of the ground truth. SUPREME was better than SUPREME-, but the difference was not significant.

Using support from the predicted subtypes by each model in SUPREME, we computed an ensembled consensus subtype based on majority voting for each patient (Supplemental File 3) and checked the survival difference between these consensus subtypes. Once again, we observed a significant (p-value=0.01) survival difference between consensus subtypes (Supplemental Figure 6C). We also observed that 882 out of 1022 patients had the same subtype prediction across all 127 models showing the robustness of SUPREME predictions.

### 2.5 Checking the impact of network-specific patient embeddings

We investigated the contribution of each patient embedding on the model performance by comparing the models built using a patient embedding from datatype 𝕏 and without using that embedding. Among all 2^𝕕^ *−* 1 models, 2^𝕕 −1^ models had the patient embedding obtained from a datatype 𝕏, called *with*𝕏_*n*_. The remaining 2^𝕕 − 1^ *−* 1 models did not have the patient embedding obtained from 𝕏, called *no*𝕏_*n*_. For each datatype 𝕏, we compared *no*𝕏_*n*_ models against *with*𝕏_*n*_ models, showing the importance of 𝕏-specific patient embedding. We did this analysis on SUPREME- (i.e., without integrating the raw features) to ensure that differences were due to the patient embeddings only.

The results on TCGA data showed that the performance of models increased or stayed the same with the inclusion of patient embeddings from all datatypes except for gene expression (Figure 4). The inclusion of EXP-based embedding showed a significant decrease in the model performance. The exclusion of CLI- and CNA-based patient embeddings had a significant drop in the model performance. Those findings agree with single model results in Section 2.3.

**Figure 4.**
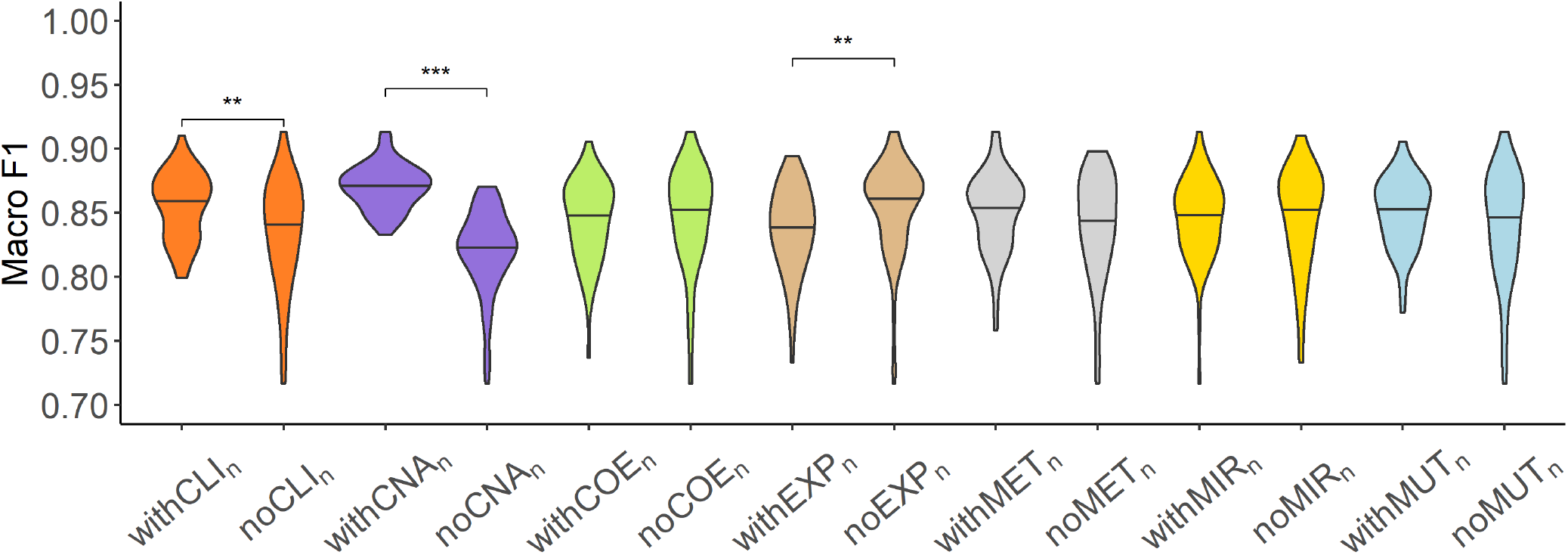
Analysis of network-specific patient embeddings. Violin plot of macro F1 scores of SUPREME-performance for the models integrated with a specific patient embedding from each datatype (*with*𝕏_*n*_ models, where 𝕏 is the datatype whose embedding is included) versus excluding that embedding (*no*𝕏_*n*_ models) on TCGA data. Significance level was measured between *with* and *no* cases of the same datatype (Wilcoxon rank-sum test p-value to compare the distribution of violin plots representing the significance *<* 0.001 by ***, else if *<* 0.01 by **, and else if *<* 0.05 by *). See Figure 1 for the abbreviations of datatypes. [SUPREME-: SUPREME without raw feature integration]

For METABRIC data, the inclusion of COE- and EXP-based embeddings increased the performance, while the other embeddings did not affect the performance much (Supplemental Figure 7A). For the combined data, MUT- and EXP-based embeddings showed higher performance when included, whereas the inclusion of CLI-based embedding did not affect the performance much (Supplemental Figure 7B).

In addition, we analyzed SUPREME results for TCGA data in terms of the best- and worst-performing models. Specifically, we had 31 top models with a macro F1 score is *≥* 0.88, and 30 bottom models with a macro F1 score is *≤* 0.83. We counted how many times each datatype occurred in the top and bottom models. CNA- and CLI-based embeddings were used for 28 and 19 out of 31 top models, respectively. The least occurred embedding was EXP-based with only six models out of 31. For the bottom models, we had 25 models from EXP-based embedding, while we had the least occurred embedding from CNA-based embedding with only five models. This analysis showed that CNA-based embedding was the most selected to have higher performance, while EXP-based embedding was the rarely selected, supporting our findings in this section and Section 2.3.

### 2.6 Checking the impact of features from each datatype

To see the impact of the features from each datatype, we ran SUPREME excluding the features from every single datatype separately. For each datatype 𝕐, we excluded 𝕐-specific node features from patient similarity networks and also did not integrate them with node embeddings during subtype prediction, called *no*𝕐_*f*_. Considering that 𝕐-specific patient similarity network was generated based on 𝕐-specific features, we compared only the combinations without 𝕐 (2^𝕕*−*1^ *−* 1 models) to ensure the differences were due to the 𝕐-specific features. We compared *no*𝕐_*f*_ models against the corresponding SUPREME models (called *with*𝕐_*f*_), to show the importance of 𝕐-specific features.

When we excluded features from any datatype, we observed a lower or comparable performance (Figure 5 and Supplemental Figure 8). The performance drop was significant for all the datatypes on TCGA, and gene expression and copy number aberration datatypes on METABRIC (Supplemental Figure 8A). The drop with the exclusion of the gene expression features was more drastic and it was consistent for all three datasets (Supplemental Figure 8B), supporting the importance of gene expression features for breast cancer (in agreement with findings in Section 2.3).

**Figure 5.**
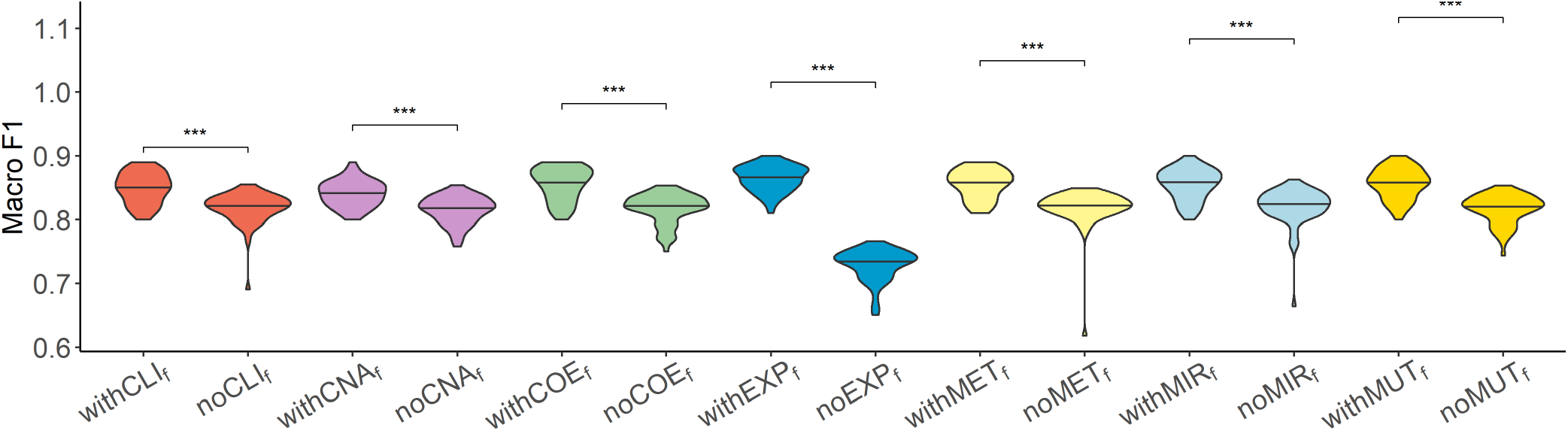
Analysis of features from each datatype. Violin plot of macro F1 scores for the models excluding the features from each datatype (*no*𝕐_*f*_ models, where 𝕐 is the datatype whose features are completely excluded) versus corresponding SUPREME models (*with*𝕐_*f*_ models) on TCGA data. Significance level was measured between *with* and *no* cases of the same datatype (Wilcoxon rank-sum test p-value to compare the distribution of violin plots representing the significance *< 0*.*001*by ***, else if *<* 0.01 by **, and else if *<* 0.05 by *). See Figure 1 for the abbreviations of datatypes

## 3 Materials and Methods

SUPREME is a computational tool for cancer subtype prediction integrating multiple biological datatypes using GCNs (Figure 1). Briefly, the first step is data preparation. In the second step, SUPREME extracts features from each datatype, and using those features, it generates individual patient similarity networks per datatype. In the third step, using the obtained networks and features, SUPREME generates the network-specific patient embeddings by running GCN on each network separately. Importantly, SUPREME utilizes local similarities generating patient embedding on each patient similarity network separately while exploiting the patient information from all the datatypes at the same time. In the last step, SUPREME does the cancer subtype prediction by integrating individual network-specific embeddings and raw multiomics features. In the following part, we explain each step of SUPREME in detail.

### 3.1 Data Preparation

To prepare the SUPREME data, we collected breast tumor samples from TCGA project data portal (available at https://www.cancer.gov/tcga) [12] and METABRIC dataset [2]. We collected all the available datatypes except for protein expression on TCGA since it was limited for breast tumor samples.

For TCGA data, we collected six datatypes, namely gene expression, microRNA expression, DNA methylation, copy number aberration, mutation, and clinical features. We preprocessed each datatype separately obtaining normalized expression values for gene expression as FPKM (fragments per kilobase of transcript per million mapped fragments), for miRNA expression as RPM (reads per million mapped reads), and obtaining gene-centric values from the segmented copy number aberration data using CNTools [29]. To eliminate redundant features, we excluded genes from the gene expression datatype unless they had expression value *≥*1 FPKM across *≥*15% of the entire cohort. Similarly, we kept microRNAs that have*≥* 0.01 RPM across *≥*30% of the entire cohort. We also applied differential expression analysis using DESeq2 [30] between normal and tumor samples for both expression datatypes to keep the cancer-related genes and microRNAs (with adjusted p-value *≤*0.01). For DNA methylation, there were two different platforms, namely HumanMethylation27 and HumanMethylation450 BeadChip, so we converted the probe-level data to gene-centric data separately for each platform, then combined the samples including common genes. To generate coexpression-based features, we ran WGCNA tool [31] using gene expression and used module eigengenes as coexpression features. For mutation data, we collected masked somatic mutation of Simple Nucleotide Variation as gene-centric across samples and converted the values to binary by modifying nonzero values to 1. We collected age, menopause status, race, tumor stage, tumor status, estrogen receptor status, progesterone receptor status, and HER2 status of samples as clinical features. Age was used as a continuous value, while other categorical values were kept as one-hot encoded. Since the number of features obtained from miRNA expression, clinical, and coexpression datatypes was *≤* 500, we did not perform any further feature selection. Due to the high dimensionality of the remaining four datatypes (i.e., gene expression, DNA methylation, copy number aberration, and mutation), we utilized a Random Forest-based feature selection algorithm Boruta [32] to keep the top features. The number of top features was a hyperparameter with the options of 200, 500, 1000, and 1500. We tuned this hyperparameter for each datatype separately during the computation of network-specific patient embeddings. Finally, all the datatypes were scaled individually by computing z-scores except for mutation features and one-hot encoded clinical features since they were binary. As the ground truth class labels, we obtained PAM50 labels of the samples. Specifically, we had five different class labels for the prediction task: Basal-like, HER2-Enriched, Luminal-A, Luminal-B, and Normal-like. Finally, including the features from coexpression, we generated features from seven datatypes in total for 1022 TCGA breast tumor samples.

For METABRIC data, we collected breast tumor samples along with four datatypes, namely gene expression (log intensity values as Illumina HT 12 v3 microarray data), copy number aberration (copy number alterations from DNA copy with calls made after correcting normal contamination and removing copy number variation), mutation (somatic mutation, Affymetrix SNP 6.0), and clinical (see [2] for data processing details). Utilizing WGCNA, coexpression features were generated as described for TCGA data. Gene expression, mutation, and copy number aberration datatypes were already preprocessed, thus we applied the same feature selection algorithm used in TCGA on them keeping the same number of features as in TCGA. We generated the same clinical features that we had in TCGA, kept mutation features binary, and collected PAM50 subtype labels. Finally, including the features from coexpression, we generated features from five datatypes in total for1699 METABRIC breast tumor samples.

In addition to these two datasets, we also generated a combined dataset using both TCGA and METABRIC through their common datatypes. To integrate gene expression from microarray (METABRIC’s gene expression) and RNA-seq (TCGA’s gene expression) platforms, we utilized FSQN tool [33], which changes the distribution of data by applying feature-specific quantile normalization, and obtained gene expression datatype. WGCNA tool requires all the samples to be processed the same way, therefore we did not have coexpression datatype. Copy number aberration data were processed differently in TCGA and METABRIC, so we did not combine this datatype, either. Combining PAM50 subtype labels, clinical features, and mutation features from both datasets, we had a total of 2721 breast tumor samples for the combined data along with three datatypes: clinical, gene expression, and mutation. The number of features and samples for each dataset are shown in Table 1.

### 3.2 Network Generation & Feature Extraction

SUPREME incorporates seven different datatypes for TCGA data, five different datatypes for METABRIC data, and three different datatypes for the combined data. The selected features in the data preprocessing step (i.e., multiomics features) were used to compute the similarity between patients when generating the patient similarity networks, as node features in the patient similarity networks, and to integrate as raw features before the prediction task. To compute patient similarities in datatype-specific patient similarity networks, we used Pearson correlation for gene expression, copy number aberration, DNA methylation, microRNA expression, and coexpression datatypes; the Gower metric [34] from the daisy function of cluster R package [35] for clinical features; and Jaccard distance for binary mutation features. The number of edges kept in the networks was a hyperparameter, which was tuned in the GCN module. After selecting the top edges, the edge weights were eliminated to generate an unweighted network. We used 2500 edges for the datatypes of TCGA, 4500 for METABRIC, and 7000 for the combined data (having approximately 2.5 times the sample size).

Patient similarities in mutation datatype were the same for a high number of patient pairs. When we need top 2500/4500/7000 edges for TCGA, METABRIC, and the combined data, we had 16,080/622,801/44,373 edges, respectively. Especially for METABRIC data, we had a high number of edges, causing high computation time. In addition to this run, we ran with randomly selected 50,000 edges and had highly similar performance with less computational time. Therefore, we used the randomly selected edges for mutation of the METABRIC dataset.

When there is a high number of raw features to integrate, this might affect the prediction performance, and training the model could be time-consuming. Thus, we added another optional feature selection step to further reduce the number of raw features integrated with the node embeddings for the prediction task. We enabled this additional feature selection for TCGA data where we had a high number of raw features and observed that it reduced running time without affecting the prediction performance.

### 3.3 Patient Embedding Generation

After extracting multiomics features and generating patient similarity networks from all the available datatypes, we generated network-specific patient embeddings, which capture the topology of the network as well as node features to be utilized in a downstream ML task.

Applying deep learning on graphs, GNN models generate the node embeddings from a given network to represent its node features and local network topology. To learn patient embeddings, we applied a GNN-based approach to utilize information from the local neighborhood of patient nodes and multiomics features associated with each node. Learning neighborhood features is achieved by an aggregation function, which varies depending on the GNN method.

In this study, we used the GCN model of Kipf and Welling [18] involving self edges into convolution and scaling the sum of aggregated features across the neighbors. GCN models learn the data by performing convolution on networks, considering one-hop local neighbors with equal contribution, and encoding the local topology of the network. Stacked layers involve recursive neighborhood diffusion considering more than a one-hop neighborhood.

Let’s call an undirected graph as *𝒢* = (*𝒱, ε*) where *𝒱* is a set of *n* nodes, i.e., *𝒱* = *{v*_1_, *v*_2_, …, *v*_*n*_*}*, and *ε* is a set of edges between nodes where (*v*_*i*_, *v*_*j*_) *∈ ε* when *v*_*i*_ *∈ 𝒱, v*_*j*_ *∈ 𝒱* and *v*_*i*_ and *v*_*j*_ have an association based on the graph *𝒢*. Since the graph *𝒢* is undirected, (*v*_*i*_, *v*_*j*_) *∈ ε ⇐⇒* (*v*_*j*_, *v*_*i*_) *∈ ε*.

The input for a GCN model is a feature matrix *𝒳 ∈* ℝ^*nxk*^ where *k* is the feature size, and the adjacency matrix *A ∈* R^*nxn*^ with added self-edges defined as:

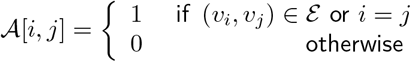

The iteration process is defined as:

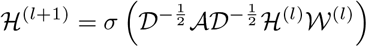

with *ℋ*(0) = *𝒳* where

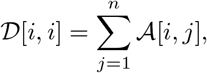

*ℋ*^(*l*)^ is the activation matrix in the *l*^*th*^ layer, 𝒲^(*l*)^ is the trainable weight matrix in the *l*^*th*^ layer and *σ* is the activation function.

Considering TCGA data, the SUPREME setup for the single model generation was as follows: there were seven single networks (i.e., the patient similarity networks explained in Section 3.2), each obtained from a different datatype. All networks had nodes as breast cancer patients and edges based on the patient similarities from the corresponding data. For instance, let us consider *𝒢* as a gene expression-derived patient similarity network. This network connects patient nodes with a high correlation between their gene expression profile. As node features, *𝒢* has the combined features, that were extracted from all the seven datatypes in the second step. Features of *v*_*i*_ is denoted as *x*_*i*_ *∈* ℝ^*k*^ where *k* is the total feature size. So, the stacked feature matrix *X ∈* ℝ^*nxk*^ is:

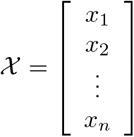

The local one-hop neighborhood of a node *v*_*i*_ is *𝒩*_*i*_ = *{v*_*j*_ : (*v*_*i*_, *v*_*j*_) *∈ ε}* that included the set of nodes having an association with the node *v*_*i*_. Feature aggregation on the local neighborhood of each node was done by multiplying *𝒳* by the *nxn*-sized scaled adjacency matrix *A′* where

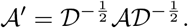

Using 2-layered GCN in SUPREME, we had the form of the forward model giving the output *Ƶ* where

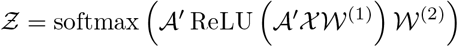

and *𝒲*^(1)^ *∈* ℝ^*kxh*^, *𝒲*^(2)^ *∈* ℝ^*hxc*^ were the trainable weights for the first and second layers, respectively, where *h* was the hidden layer size and *c* was the number of classes to predict (namely, Basal-like, Luminal-A, Luminal-B, HER2-Enriched, and Normal-like, with *c* = 5). The loss function was calculated by cross-entropy error. Adam optimization [36] was used as the state-of-the-art for stochastic gradient descent algorithm and dropout was added for the first GCN layer. Early stopping was used with the patience of 30 forced to have at least 200 epochs

We split 20% of the total samples as a test set and the test set was kept for the final evaluation of the tool. This splitting was stratified, that is, keeping the same ratio of the subtype labels in the original data for each split. The remaining 80% of the samples were used for training (60%) and validation (20%), and those splits were randomly selected for each run as stratified. To tune the hyperparameters of the GCN model (i.e., hidden layer size and learning rate), for each run, SUPREME repeated an evaluation metric (i.e., macro F1 score) 10 times for each hyperparameter combination (See Supplemental File 2) and selected the hyperparameter combination giving the best median macro F1 score on the validation data to generate the final model. These final models were used to extract network-specific patient embeddings to use in the prediction task.

Similarly applying the methodology for other datatypes, we generated seven different GCN models on TCGA data. Repeating the same procedure for other datasets, we obtained five models on the METABRIC data and three models on the combined data.

### 3.4 Cancer Subtype Prediction

We concatenated *all the different combinations* of patient embeddings (having separate 2^𝕕^*−* 1 prediction models), integrated them with raw features, and evaluated the models. Specifically, we had 127, 31, and seven different SUPREME models for TCGA, METABRIC, and the combined data, respectively.

We tested SUPREME with several ML methods namely, XGBoost, SVM, RF, and MLP algorithms. For all datasets, we decided to use MLP as it gave consistently high performance (Supplemental Table 1 and discussion section for the details). We did hyperparameter tuning for the prediction task, similar to GCN hyperparameter tuning in the previous step. We used the training and validation cohort to tune the hyperparameters (e.g., hidden layer size and learning rate) of the final model, where training and validation splits were randomly selected as stratified. We repeated the SUPREME run 10 times for each hyperparameter combination and used the hyperparameter combination giving the best median macro F1 score on the validation data. Using this hyperparameter combination, the final model was built and evaluated 10 times on the test data, which was never seen during training and hyperparameter tuning. The evaluation metrics (macro F1 score, weighted F1 score, and accuracy) were obtained from the median of these 10 runs.

### 3.5 Supervised Tool Comparison

We compared SUPREME with cancer (sub)type prediction tools and baseline methods, namely MOGONET [27], GCNC [26], DeepCC [28], MLP, RF, SVM, and XGBoost. Similar to SUPREME, we ran all the possible combinations for the integrative tools and ML-based baseline methods excluding DeepCC and GCNC, which cannot run on multiple datatypes. For GCNC, we generated a coexpression network and protein-protein interaction network following the original paper and evaluated two prediction models using only the gene expression datatype. For DeepCC, we built the model using only the gene expression datatype, too. For ML-based baseline methods (i.e., MLP, RF, SVM, and XGBoost), we integrated only the raw features from the selected combination and did the prediction with those features. Even though MOGONET is applicable to any number of datatypes, we could not run the tool for the models with more than five datatypes (waiting time was more than two days per combination), thus we had only 31 different models for TCGA data, while we had all models for METABRIC and the combined data.

### 3.6 Survival Analysis

To measure the power of methods in differentiating groups with survival differences, we did a survival analysis using the log-rank test to obtain a p-value to measure the difference between survival curves of different cancer subtype cohorts. From TCGA data, patient survival-related information (days to death, days to the last follow-up, and vital status) were collected. Patients with “dead” vital status were considered dead, otherwise censored.

We did survival analysis on the clusters obtained from popular unsupervised cancer subtyping methods, namely iClusterPlus [13], SNF [14], and PINSPlus [15]. We gave all the different combinations of selected features of datatypes as input, obtained five clusters consistent with PAM50 subtypes, and compared the survival differences between them. In addition to the unsupervised methods, we did survival analysis for the integrative supervised tools (i.e., the baseline methods and the GCN-based approaches). For the supervised tools, we defined the groups based on the predicted subtypes and checked the survival differences between them.

SNF generates multiple networks from multiple datatypes and does spectral clustering on the fused network. For single datatypes, since SNF is unable to create a fused network, we externally applied spectral clustering. In addition, SNF did not converge for some combinations as reported in other studies [37]. We ignored those models in our comparison. For iClusterPlus, the tool itself can handle a maximum of four datatypes, so we could not run the combinations with more than four datatypes, having at most 15 models per dataset. We generated all possible combinations for the other tools to check their overall performance.

### 3.7 Ablation Study

We compared our tool with its variations when some steps were skipped to assess their importance (Table 3). To check the performance of different methods, we compare them with all the combinations of datatypes (Section 2.2). In this section, a comparison of our method with SUPREME-showed the importance of raw feature integration. Also, to show the importance of GCN-based approaches, we trained the same ML algorithm (MLP in our case) using only the raw features and compared it with the SUPREME, which was based on the same raw features and additional patient embeddings.

To show the impact of each datatype separately, we demonstrated the performance of SUPREME models based on a single data type (Section 2.3). We also compared SUPREME with other methods that can work with a single data modality only. To show the importance of embeddings at a single datatype level, we compared SUPREME with the MLP model trained on the features from the corresponding datatype.

To see the importance of each network-specific embedding, we checked the performance of patient embedding combinations including one datatype (called with𝕏_n_, where 𝕏 was the datatype whose embeddings were included) versus excluding that datatype (called no𝕏_*n*_, where 𝕏 was the datatype whose embeddings were excluded) (Section 2.5).

To demonstrate the effect of features from each datatype, we excluded those features completely (Section 2.6). To ensure complete exclusion of the features of a datatype, we did not use these features as node features in all patient similarity networks, we excluded them during raw feature integration, and we did not use the patient similarity network generated based on these features. The exclusion of a network resulted in the exclusion of its network-specific patient embedding, so we had 2^𝕕*−*1^ *−* 1 models out of 2^𝕕^ *−* 1. We called these models no𝕐_*f*_ models, where 𝕐 was the datatype whose features were excluded as specified. We compared these models with the corresponding 2^𝕕*−*1^ *−* 1 original models (called with𝕐_*f*_ models).

## 4 Discussion

We introduced SUPREME, a novel integrative approach utilizing GCNs on multiple patient similarity networks where patient nodes are attributed with multiomics features. We made SUPREME a publicly available tool for researchers, biologists, and clinicians to utilize, available at https://github.com/bozdaglab/SUPREME under Creative Commons Attribution Non Commercial 4.0 International Public License. Running SUPREME on three datasets, we obtained models for all combinations of 𝕕 datatypes (i.e., 2^𝕕*−* 1^ models). We compared SUPREME with seven cancer (sub)type prediction tools and baseline methods and observed that SUPREME substantially outperformed or was on par with them based on overall macro F1 score, accuracy, and weighted F1 score (Figure 2, Supplemental Table 2, and Supplemental Figures 3 and 4). We differentiated Normal-like subtype, which has the smallest sample size for three datasets, significantly better than all other tools on all three datasets showing SUPREME’s robustness even for minority classes (Supplemental Figure 5).

We applied survival analysis to see the power of the methods to differentiate subtypes having significant survival differences. Using TCGA data, we compared our tool with nine popular integrative cancer subtype differentiating tools and baseline methods and SUPREME had consistently significant survival differences between predicted subtypes outperforming the other tools (Figure 3).

Based on the majority of predictions, we determined ensemble subtype labels, most of which had high support from individual models (Supplemental File 3). We observed that survival difference between these ensemble subtypes was more significant than survival difference between gene expression-based ground truth (i.e., PAM50) subtypes (Supplemental Figure 6). These results suggest that some survival-related characteristics cannot be explained by gene expression data alone. SUPREME was able to extract these survival-related characteristics utilizing additional data modalities. SUPREME’s ensemble label predictions that were different from ground truth with high support could be further examined by biologists and clinicians.

To show the effect of main steps of SUPREME, we performed an ablation study. In addition, we analyzed datatype-specific embeddings and datatype-specific features. We found that gene expression features were highly important for single models and overall, as expected for breast cancer. Findings about the important embeddings of datasets in Sections 2.5 were supported by SUPREME-single models in Section 2.3, where models were fed by only one embedding. We observed that patient embeddings were mostly more informative than raw features. Integrating raw features with patient embeddings usually improved the model performance (Figure 2, Supplementary Figure 3) except for raw features from few datatypes in single datatype-based models (Table 2, and Supplemental Tables 3, 4 and 5).

To compare the performance when we do not utilize the local neighborhood, we ran SUPREME- on TCGA data with the EXP-based single model when we do not have any neighbor than the patient itself. In that model, we had a macro F1 score of 0.85±0.02 for SUPREME-, which was much higher than the original EXP-based model of SUPREME-. This model was even better than the EXP-based single model of SUPREME. This might suggest that EXP-based patient features themselves could perform better than neighborhood-convolved features because the ground truth utilizes patient features themselves to decide the subtype labels. Similarly, because of that, we might see a performance improvement when we add EXP-based raw features.

SUPREME provides four options of ML algorithms to integrate embeddings and raw features, namely MLP, RF, SVM, and XGBoost. We ran SUPREME with all these choices and compared the performances (Supplemental Table 1). RF and XGBoost had a low performance for some models. Overall, SVM had a good performance on every three datasets, however, it did not converge for some models. For this study, we chose MLP due to its high and consistent prediction performance for all three datasets and its low running time.

In our experiments, we observed a high number of edges in mutation-based patient similarity networks as there were many patient pairs with the same similarity. Furthermore, the MUT-based models on TCGA data had a high predictive performance, whereas these models had a low predictive performance on METABRIC and the combined datasets. These discrepancies were mainly due to the sparse nature of the binary mutation feature. For the special datatypes with binary-like sparse values like mutation, patient similarity networks and extracted features could be generated in a more sophisticated way such as based on the functional effect of these mutations [38, 39, 40, 41].

SUPREME is extendable to any number of datatypes to integrate. For cases where many datasets are integrated, to avoid potential overfitting, SUPREME provides an optional feature selection step for raw features before training the final prediction model. Users could skip raw feature integration altogether when network-specific patient embeddings provide sufficient discriminatory power. Users could run SUPREME on their training/validation data by enabling/disabling these features to optimize their models. In addition, users could perform ablation studies on SUPREME to determine the most effective data modalities and their combinations. Depending on these results, for the final prediction, users could rely on the most effective model or an ensemble model utilizing the most promising features and networks.

As a future direction, SUPREME could utilize attention mechanisms [42, 43, 44], which allows getting weighted contribution from different datatypes, and also weighted neighborhood from networks. In addition to multiomics datatypes, there are some regulatory relations such as competing endogenous RNA (ceRNA) regulation, which has been recently discovered with important insights into cancer [45]. In our recent work, we inferred ceRNA interactions in breast cancer [46]. To adopt this kind of regulatory relations, SUPREME could be improved to utilize patient similarity networks based on gene regulatory interactions and more complex patient relations. Improving the existing methodologies with recent advances in the literature, we can obtain more clear cancer subtype groups to pave the way for precision medicine.

## Acknowledgements

This work was supported by the National Institute of General Medical Sciences of the National Institutes of Health under Award Number R35GM133657.

