## Supplemental Tables and Figures for "SUPREME: A cancer subtype prediction methodology integrating multiomics data using Graph Convolutional Neural Network"

### 1 Supplementary Tables

**Table 1. SUPREME performance with different integration methods** Minimum, median, and maximum macro F1 of all the different models across different integration methods. [SUPREME-: SUPREME without raw feature integration]

| Setup | TCGA | METABRIC | Combined |
|---|---|---|---|
| SUPREME (SUPREME$^{MLP}$) | **0.80**, 0.86, 0.90 | **0.80, 0.82**, 0.84 | **0.79, 0.80**, 0.81 |
| SUPREME- (SUPREME-$^{MLP}$) | 0.72, 0.85, 0.91 | 0.34, 0.78, 0.80 | 0.63, 0.77, 0.79 |
| SUPREME$^{RF}$ | 0.14, 0.84, 0.91 | 0.11, 0.79, 0.82 | 0.75, 0.79, 0.79 |
| SUPREME-$^{RF}$ | 0.14, 0.81, 0.92 | 0.35, 0.79, 0.82 | 0.59, 0.73, 0.79 |
| SUPREME$^{SVM}$ | **0.80, 0.87**, 0.92 | 0.79, **0.82, 0.85** | 0.78, **0.80**, 0.81 |
| SUPREME-$^{SVM}$ | 0.71, **0.87, 0.94** | 0.37, 0.79, 0.82 | 0.76, 0.78, 0.80 |
| SUPREME$^{XGBoost}$ | 0.74, 0.84, 0.92 | 0.76, 0.79, 0.83 | 0.78, **0.80, 0.83** |
| SUPREME-$^{XGBoost}$ | 0.67, 0.85, 0.93 | 0.35, 0.78, 0.84 | 0.62, 0.77, 0.79 |

**Table 2. Macro F1 score, weighted F1 score and accuracy for the supervised tools that use only gene expression datatype.**

**Macro F1 scores**

| Method | TCGA | METABRIC | Combined |
|---|---|---|---|
| DeepCC | 0.81±0.02 | 0.79±0.02 | 0.82±0.01 |
| $GCNC_{PPI}$ | 0.81±0.02 | 0.80±0.02 | 0.81±0.02 |
| $GCNC_{COE}$ | 0.82±0.03 | 0.80±0.02 | 0.80±0.02 |

**Weighted F1 scores**

| Method | TCGA | METABRIC | Combined |
|---|---|---|---|
| DeepCC | 0.90±0.01 | 0.83±0.02 | 0.85±0.01 |
| $GCNC_{PPI}$ | 0.88±0.01 | 0.83±0.02 | 0.84±0.01 |
| $GCNC_{COE}$ | 0.88±0.01 | 0.82±0.02 | 0.84±0.01 |

**Accuracies**

| Method | TCGA | METABRIC | Combined |
|---|---|---|---|
| DeepCC | 0.90±0.01 | 0.83±0.02 | 0.86±0.01 |
| $GCNC_{PPI}$ | 0.88±0.01 | 0.84±0.02 | 0.84±0.01 |
| $GCNC_{COE}$ | 0.88±0.01 | 0.83±0.02 | 0.84±0.01 |

**Table 3. Single model results on TCGA data** Weighted F1 score and accuracy for each model with a single dataype. [Datatypes are abbreviated as CLI: clinical, CNA: copy number aberration, COE: coexpression, EXP: gene expression, MET: DNA Methylation, MIR: microRNA expression, MUT: mutation. MLP stands for Multi-layer Perceptron, SUPREME-: SUPREME without raw feature integration.]

**Weighted F1 scores**

| Method | CLI | CNA | COE | EXP | MET | MIR | MUT |
|---|---|---|---|---|---|---|---|
| SUPREME | 0.80±0.02 | **0.87±0.02** | 0.85±0.01 | **0.89±0.01** | **0.87±0.02** | **0.83±0.01** | **0.83±0.01** |
| SUPREME- | **0.83±0.01** | 0.86±0.01 | **0.86±0.01** | 0.86±0.02 | **0.87±0.02** | 0.82±0.01 | **0.83±0.01** |
| MLP | 0.62±0.04 | 0.68±0.03 | 0.77±0.01 | 0.88±0.01 | 0.79±0.02 | **0.83±0.02** | 0.49±0.06 |
| MOGONET | 0.58±0.01 | 0.68±0.01 | 0.75±0.01 | 0.82±0.00 | 0.73±0.01 | 0.78±0.02 | 0.53±0.00 |

**Accuracies**

| Method | CLI | CNA | COE | EXP | MET | MIR | MUT |
|---|---|---|---|---|---|---|---|
| SUPREME | 0.81±0.01 | **0.87±0.02** | **0.86±0.01** | **0.89±0.01** | **0.88±0.02** | 0.83±0.01 | **0.83±0.01** |
| SUPREME- | **0.83±0.02** | 0.**87±0.01** | **0.86±0.01** | 0.87±0.01 | 0.87±0.02 | 0.82±0.01 | **0.83±0.01** |
| MLP | 0.71±0.03 | 0.70±0.03 | 0.79±0.01 | 0.88±0.01 | 0.80±0.02 | **0.84±0.02** | 0.55±0.03 |
| MOGONET | 0.62±0.02 | 0.70±0.01 | 0.77±0.01 | 0.82±0.00 | 0.74±0.00 | 0.78±0.02 | 0.56±0.01 |

**Table 4. Single model results on METABRIC data** Macro F1 score, Weighted F1 score, and accuracy for each model with a single dataype. [Datatypes are abbreviated as CLI: clinical, CNA: copy number aberration, COE: coexpression, EXP: gene expression, MUT: mutation. MLP stands for Multi-layer Perceptron, SUPREME-: SUPREME without raw feature integration.]

**Macro F1 scores**

| Method | CLI | CNA | COE | EXP | MUT |
|---|---|---|---|---|---|
| SUPREME | **0.49±0.01** | 0.45±0.02 | **0.75±0.01** | 0.78±0.02 | **0.36±0.03** |
| SUPREME- | **0.49±0.03** | 0.44±0.01 | **0.75±0.01** | **0.82±0.01** | 0.33±0.04 |
| MLP | 0.46±0.02 | **0.46±0.02** | 0.71±0.02 | 0.81±0.02 | 0.30±0.03 |
| MOGONET | 0.41±0.02 | 0.42±0.01 | 0.65±0.03 | 0.73±0.01 | 0.22±0.00 |

**Weighted F1 scores**

| Method | CLI | CNA | COE | EXP | MUT |
|---|---|---|---|---|---|
| SUPREME | **0.54±0.01** | 0.52±0.02 | **0.78±0.01** | 0.81±0.02 | **0.45±0.03** |
| SUPREME- | **0.54±0.04** | 0.51±0.01 | **0.78±0.01** | **0.84±0.01** | 0.41±0.04 |
| MLP | 0.50±0.03 | **0.53±0.01** | 0.76±0.02 | **0.84±0.01** | 0.41±0.04 |
| MOGONET | 0.47±0.01 | 0.50±0.01 | 0.69±0.01 | 0.77±0.01 | 0.31±0.00 |

**Accuracies**

| Method | CLI | CNA | COE | EXP | MUT |
|---|---|---|---|---|---|
| SUPREME | **0.58±0.01** | **0.56±0.02** | **0.79±0.01** | 0.82±0.02 | **0.49±0.01** |
| SUPREME- | **0.58±0.01** | 0.54±0.01 | 0.78±0.01 | 0.84±0.01 | **0.49±0.01** |
| MLP | **0.58±0.01** | **0.56±0.01** | 0.75±0.02 | **0.85±0.01** | 0.47±0.01 |
| MOGONET | 0.52±0.02 | 0.52±0.01 | 0.70±0.01 | 0.77±0.01 | 0.46±0.00 |

**Table 5. Single model results on the combined data (TCGA + METABRIC)** Macro F1 score, Weighted F1 score, and accuracy for each model with a single dataype. [Datatypes are abbreviated as CLI: clinical, EXP: gene expression, MUT: mutation. MLP stands for Multi-layer Perceptron, SUPREME-: SUPREME without raw feature integration.]

**Macro F1 scores**

| **Method** | **CLI** | **EXP** | **MUT** |
|---|---|---|---|
| SUPREME | 0.43±0.01 | **0.81±0.01** | **0.33±0.03** |
| SUPREME- | 0.43±0.01 | 0.80±0.01 | **0.33±0.01** |
| MLP | **0.44±0.01** | 0.80±0.01 | 0.31±0.03 |
| MOGONET | 0.41±0.00 | 0.71±0.01 | 0.32±0.01 |

**Weighted F1 scores**

| **Method** | **CLI** | **EXP** | **MUT** |
|---|---|---|---|
| SUPREME | 0.53±0.01 | **0.85±0.01** | **0.47±0.03** |
| SUPREME- | 0.52±0.02 | 0.84±0.01 | 0.46±0.01 |
| MLP | **0.54±0.01** | 0.84±0.01 | 0.43±0.02 |
| MOGONET | 0.50±0.00 | 0.77±0.01 | 0.46±0.01 |

**Accuracies**

| **Method** | **CLI** | **EXP** | **MUT** |
|---|---|---|---|
| SUPREME | **0.58±0.00** | **0.85±0.01** | **0.51±0.02** |
| SUPREME- | **0.58±0.01** | 0.84±0.01 | **0.51±0.01** |
| MLP | **0.58±0.01** | **0.85±0.01** | 0.48±0.02 |
| MOGONET | 0.55±0.00 | 0.78±0.01 | 0.48±0.01 |

#### 2 Supplementary Figures

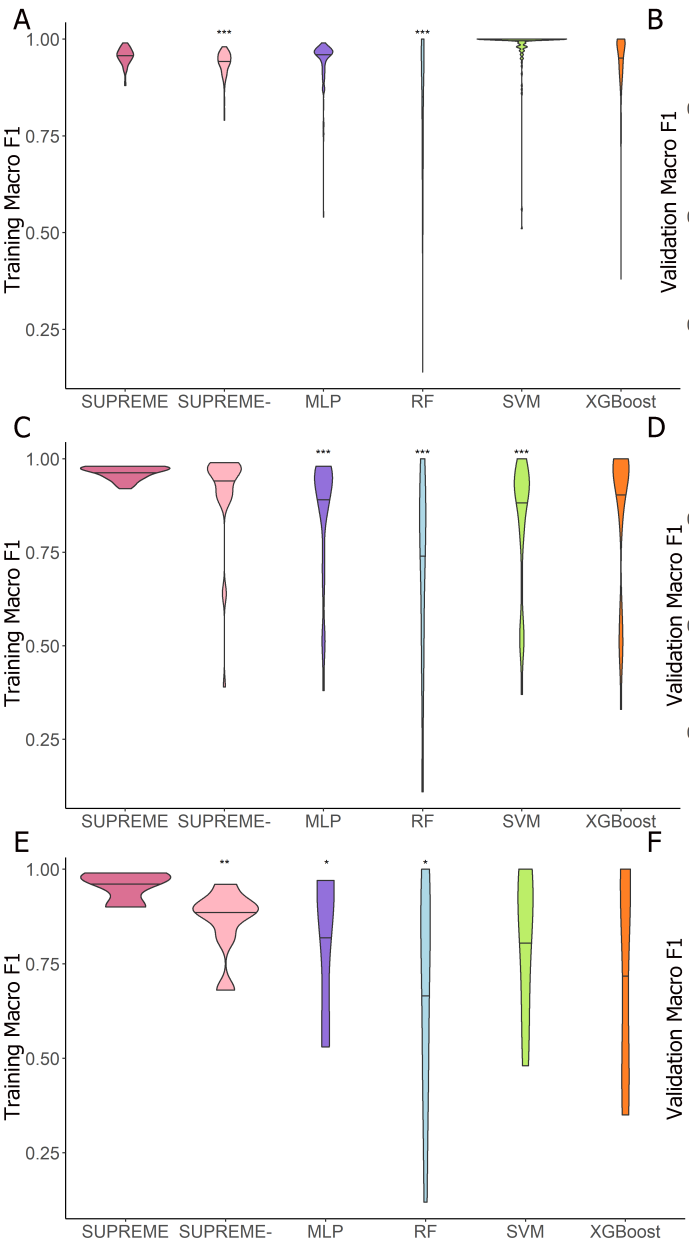


**Figure 1. Training/Validation results.** Violin plot of macro F1 scores for training and validation for SUPREME and other supervised tools on **(A-B)** TCGA data, **(C-D)** METABRIC data and **(E-F)** the combined data. The significance level was measured with respect to SUPREME (Wilcoxon rank-sum test p-value to compare the distribution of violin plots representing the significance $< 0.001$ by ***, else if $< 0.01$ by **, and else if $< 0.05$ by *). [MLP: Multi-layer Perceptron, RF: Random Forest, SUPREME-: SUPREME without raw feature integration, SVM: Support Vector Machine]

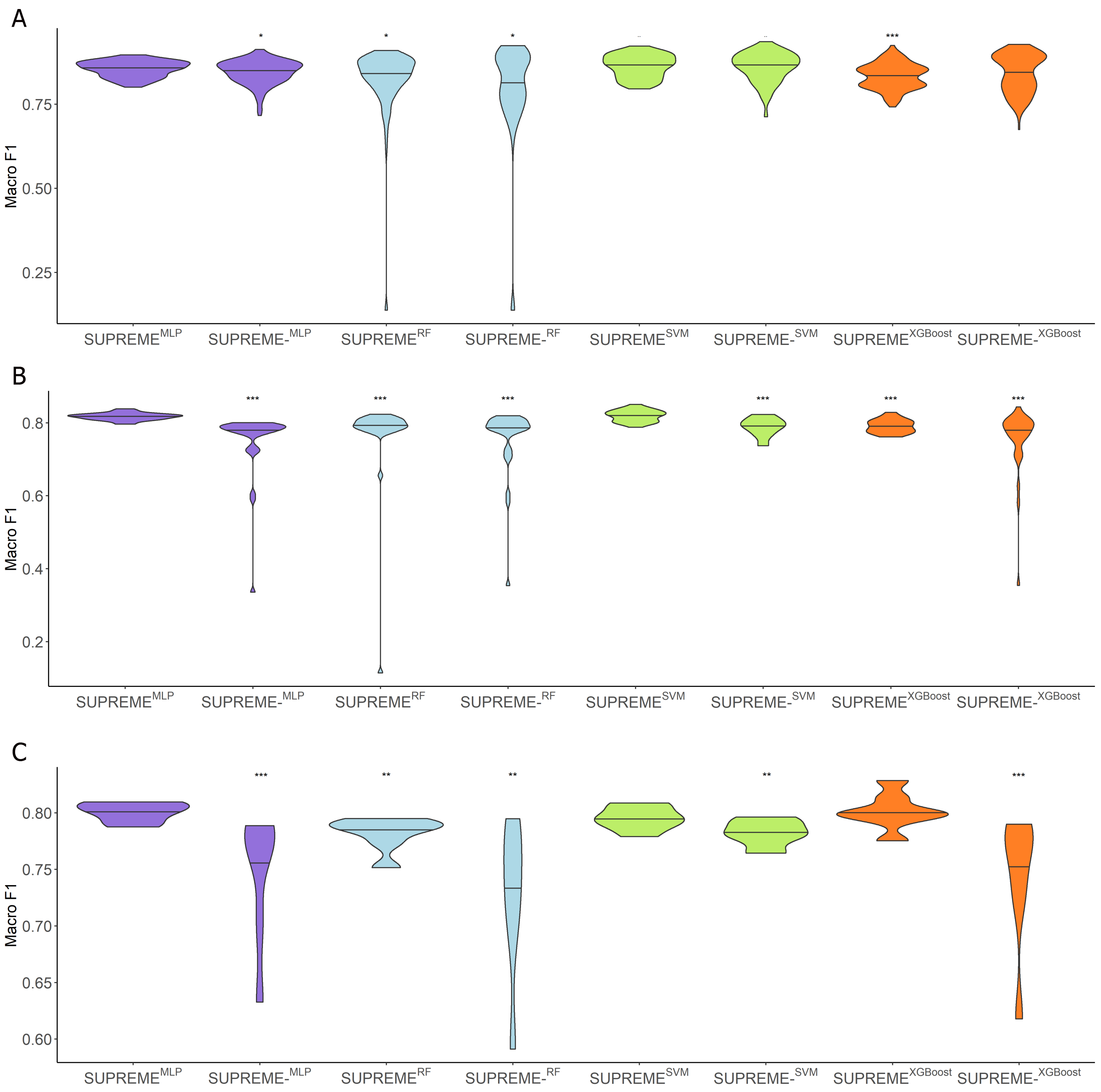


**Figure 2. Results with different integration methods.** Violin plot of macro F1s with different integration methods on **(A)** TCGA data, **(B)** METABRIC data and **(C)** the combined data. The significance level was measured with respect to SUPREME$^{MLP}$. (Wilcoxon rank-sum test p-value to compare the distribution of violin plots representing the significance $< 0.001$ by ***, else if $< 0.01$ by **, and else if $< 0.05$ by *.) [MLP: Multi-layer Perceptron, RF: Random Forest, SUPREME-: SUPREME without raw feature integration, SVM: Support Vector Machine]

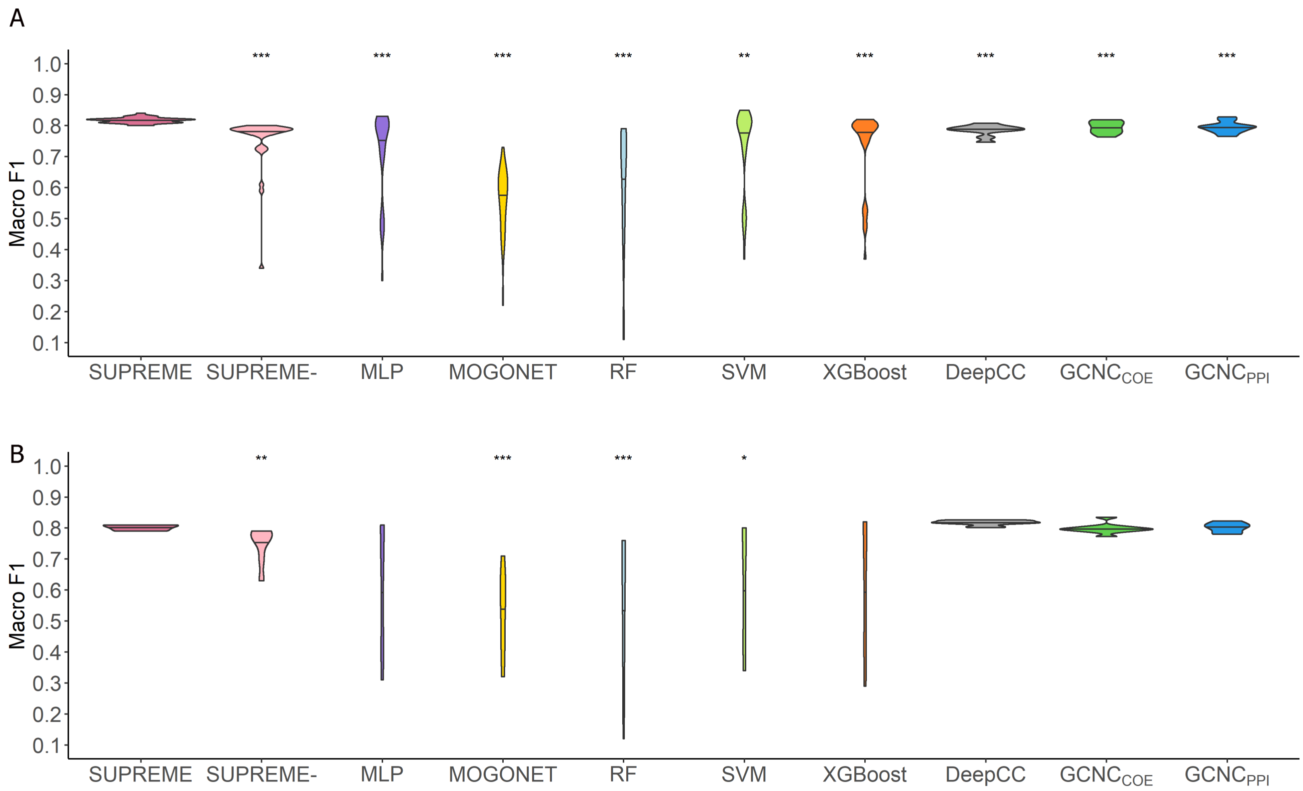


**Figure 3. Classification results.** Violin plot of macro F1 scores obtained from all different combinations of datatypes as compared to the cancer subtype prediction tools and baseline supervised methods on **(A)** METABRIC data and **(B)** the combined data. DeepCC and GCNC violin plots show the distribution of macro F1 scores of ten runs of a single model as they can only utilize gene expression datatype. The significance level was measured with respect to SUPREME (Wilcoxon rank-sum test p-value to compare the distribution of violin plots representing the significance $< 0.001$ by ***, else if $< 0.01$ by **, and else if $< 0.05$ by *). [MLP: Multi-layer Perceptron, RF: Random Forest, SUPREME-: SUPREME without raw feature integration, SVM: Support Vector Machine]

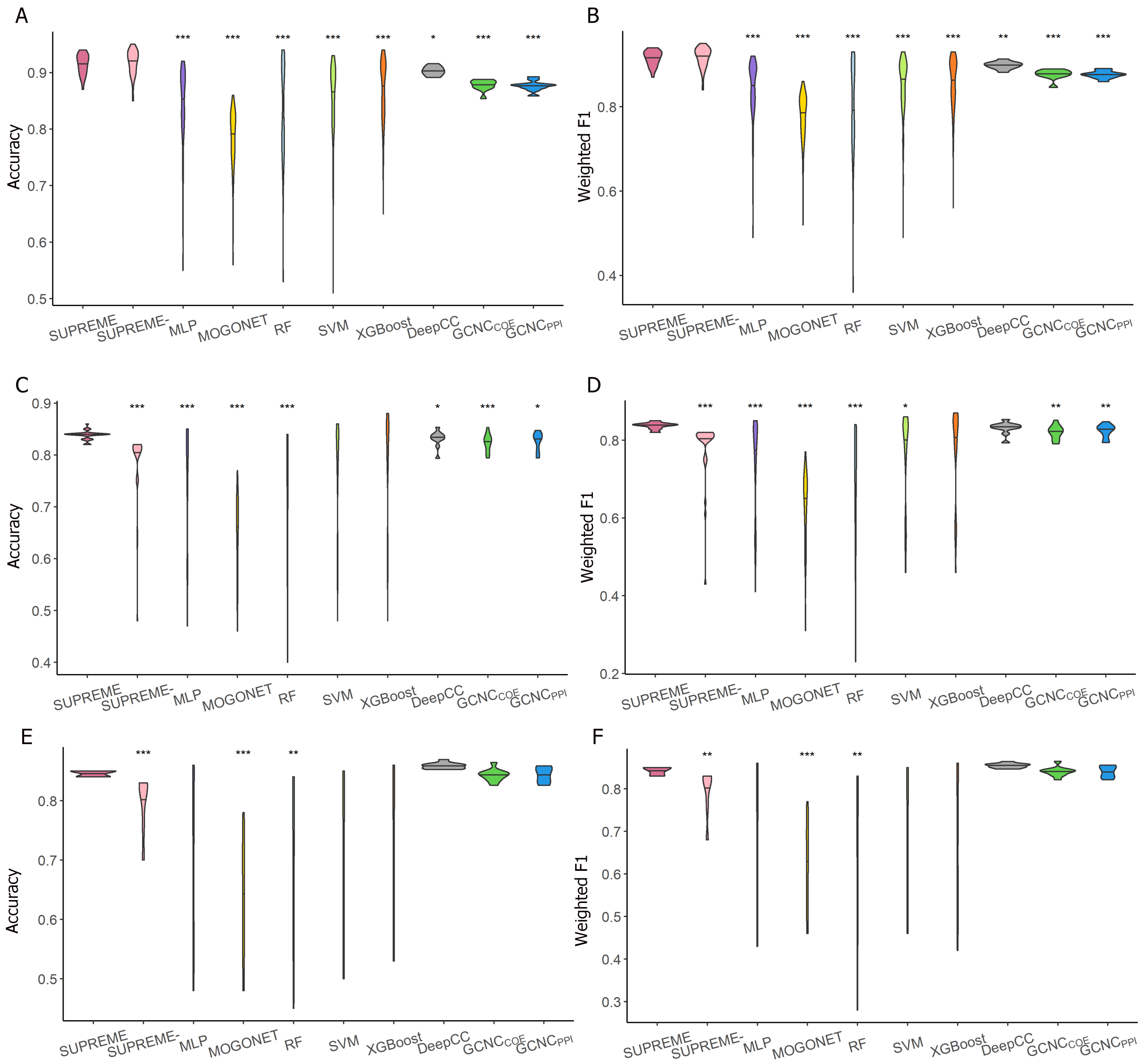


**Figure 4. Classification results as accuracy and weighted F1.** Violin plot of accuracies and weighted F1 scores obtained from all different combinations of datatypes as compared to the cancer subtype prediction tools and baseline supervised methods. **(A)** accuracy on TCGA data, **(B)** weighted F1 score on TCGA data, **(C)** accuracy on METABRIC data, **(D)** weighted F1 score on METABRIC data, **(E)** accuracy on the combined data, and **(F)** weighted F1 score on the combined data. DeepCC and GCNC violin plots show the distribution of evaluation metrics of ten runs of a single model as they can only utilize gene expression datatype. The significance level was measured with respect to SUPREME (Wilcoxon rank-sum test p-value to compare the distribution of violin plots representing the significance $< 0.001$ by ***, else if $< 0.01$ by **, and else if $< 0.05$ by *). [MLP: Multi-layer Perceptron, RF: Random Forest, SUPREME-: SUPREME without raw feature integration, SVM: Support Vector Machine]

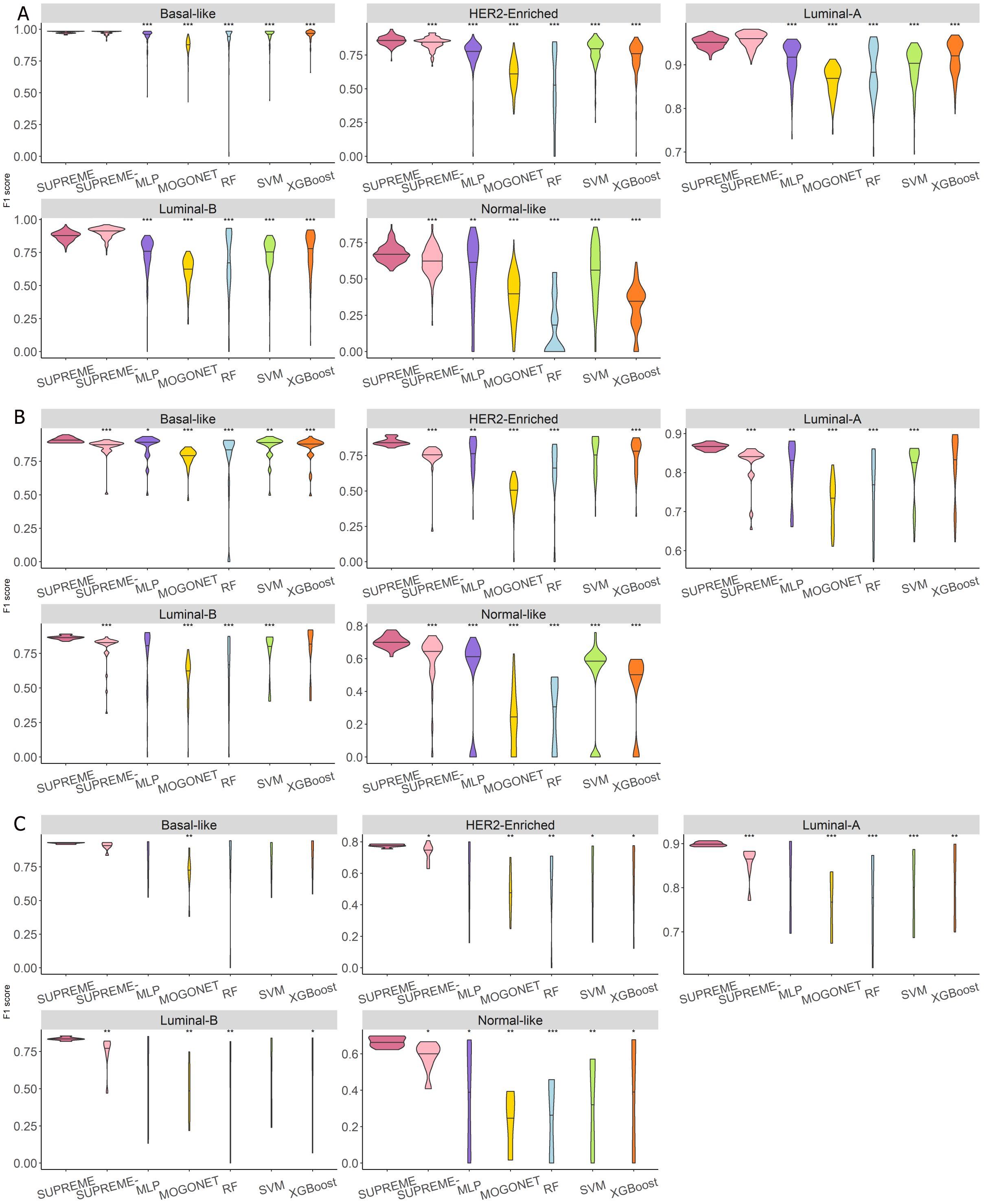


**Figure 5. Subtype-specific F1 scores.** Violin plot of F1 scores obtained from all different combinations of datatypes as compared to the cancer subtype prediction tools and baseline supervised methods on **(A)** TCGA data, **(B)** METABRIC data, and **(C)** the combined data. Significance level was measured with respect to SUPREME. (Wilcoxon rank-sum test p-value to compare the distribution of violin plots representing the significance $< 0.001$ by ***, else if $< 0.01$ by **, and else if $< 0.05$ by *.) [MLP: Multi-layer Perceptron, RF: Random Forest, SUPREME-: SUPREME without raw feature integration, SVM: Support Vector Machine]

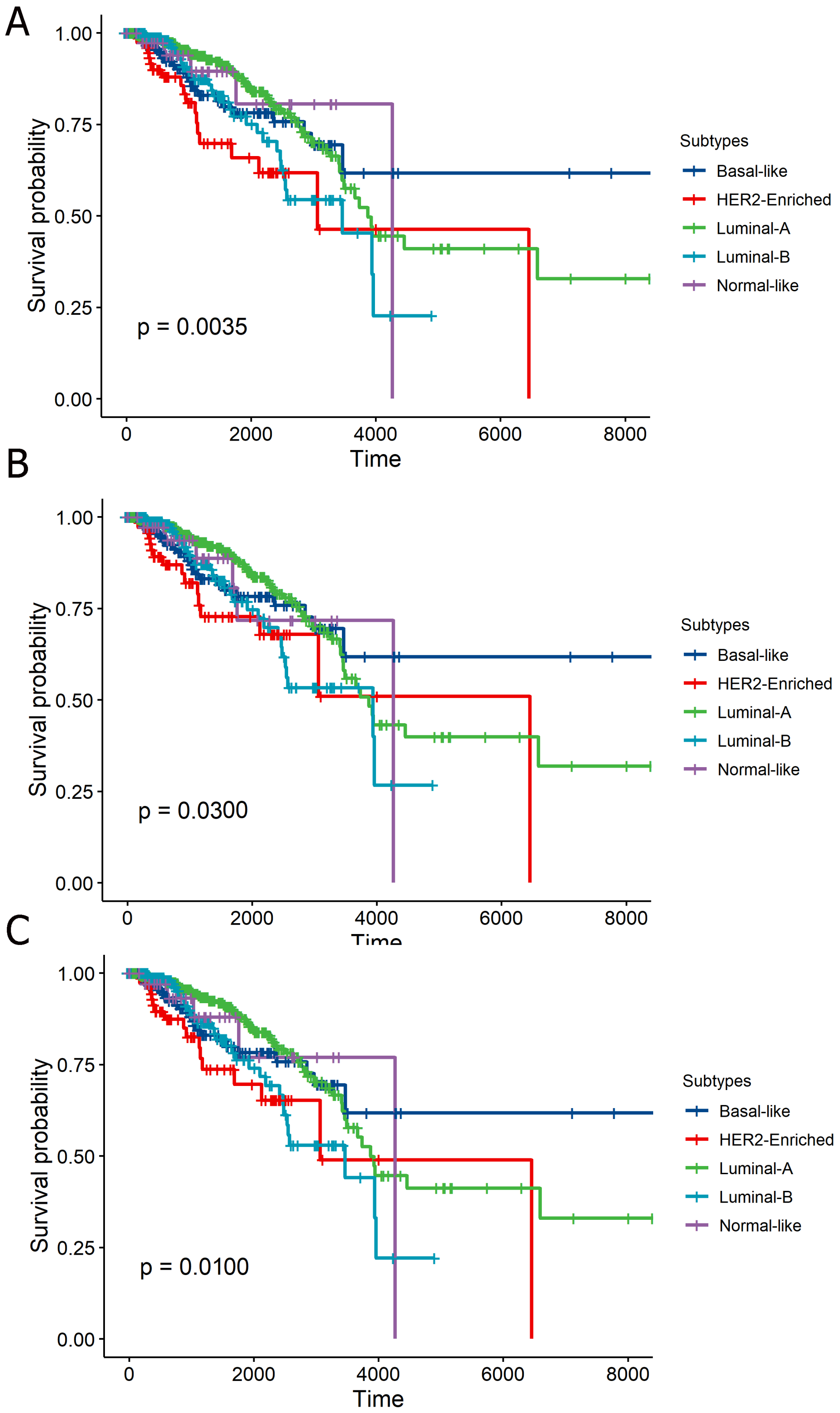


**Figure 6. Kaplan-Meier plots.** Kaplan-Meier plots of three cases: **(A)** SUPREME with the most significant survival difference (when integrating copy number aberration, coexpression, DNA methylation, and mutation-based patient embeddings), **(B)** PAM50 labels used as ground truth, and **(C)** SUPREME with ensembled labels.

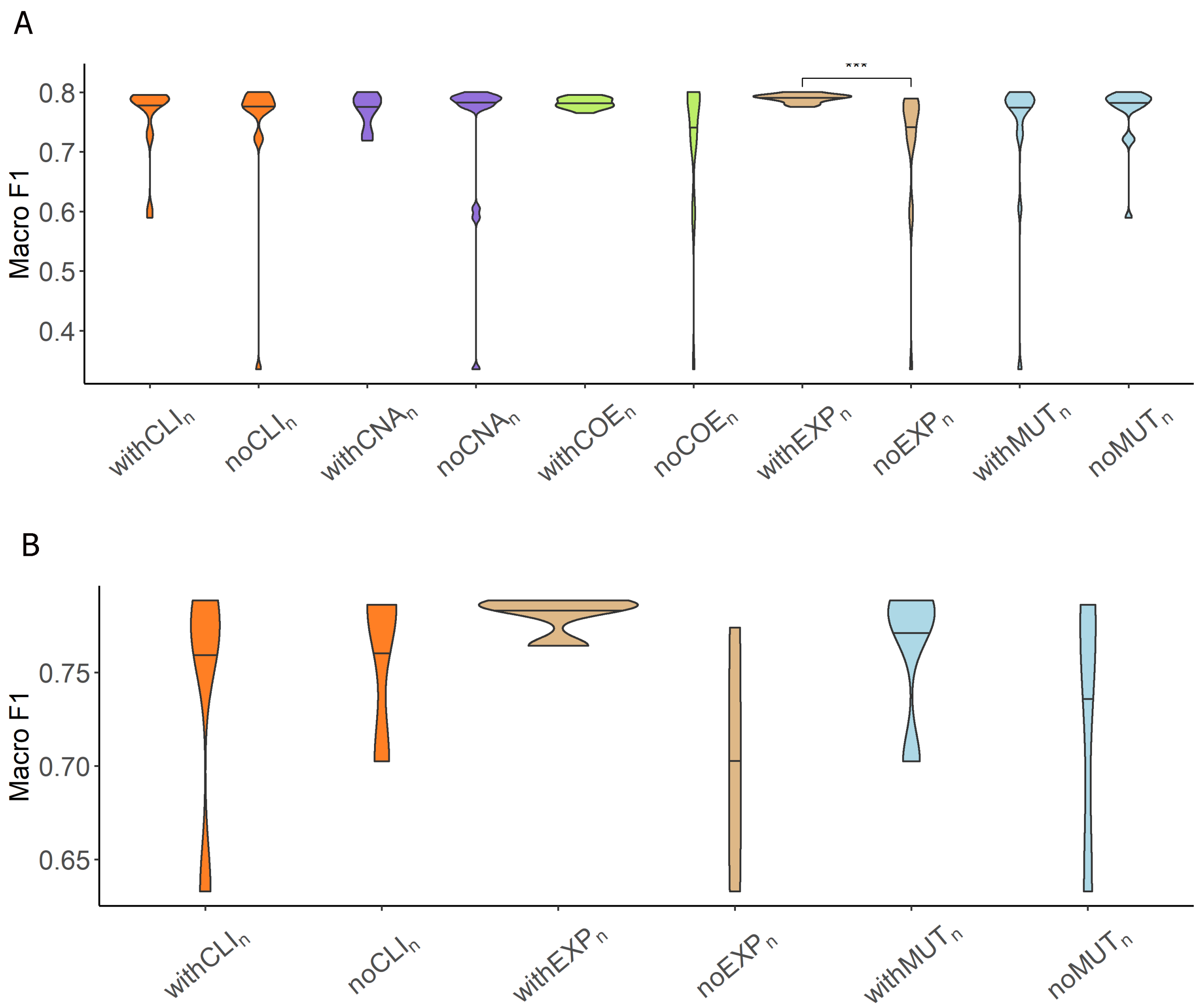


**Figure 7. Analysis of network-specific patient embeddings** Violin plot of macro F1 scores of SUPREME-performance for the models integrated with a specific patient embedding from each datatype (*with*$\mathbb{X}_n$ models, where $\mathbb{X}$ is the datatype whose embedding is included) versus excluding that embedding (*no*$\mathbb{X}_n$ models) on **(A)** METABRIC data and **(B)** the combined data. Significance level was measured between *with* and *no* cases of the same datatype (Wilcoxon rank-sum test p-value to compare the distribution of violin plots representing the significance $< 0.001$ by ***, else if $< 0.01$ by **, and else if $< 0.05$ by *). [Datatypes are abbreviated as CLI: clinical, CNA: copy number aberration, COE: coexpression, EXP: gene expression, MUT: mutation.] [SUPREME-: SUPREME without raw feature integration]

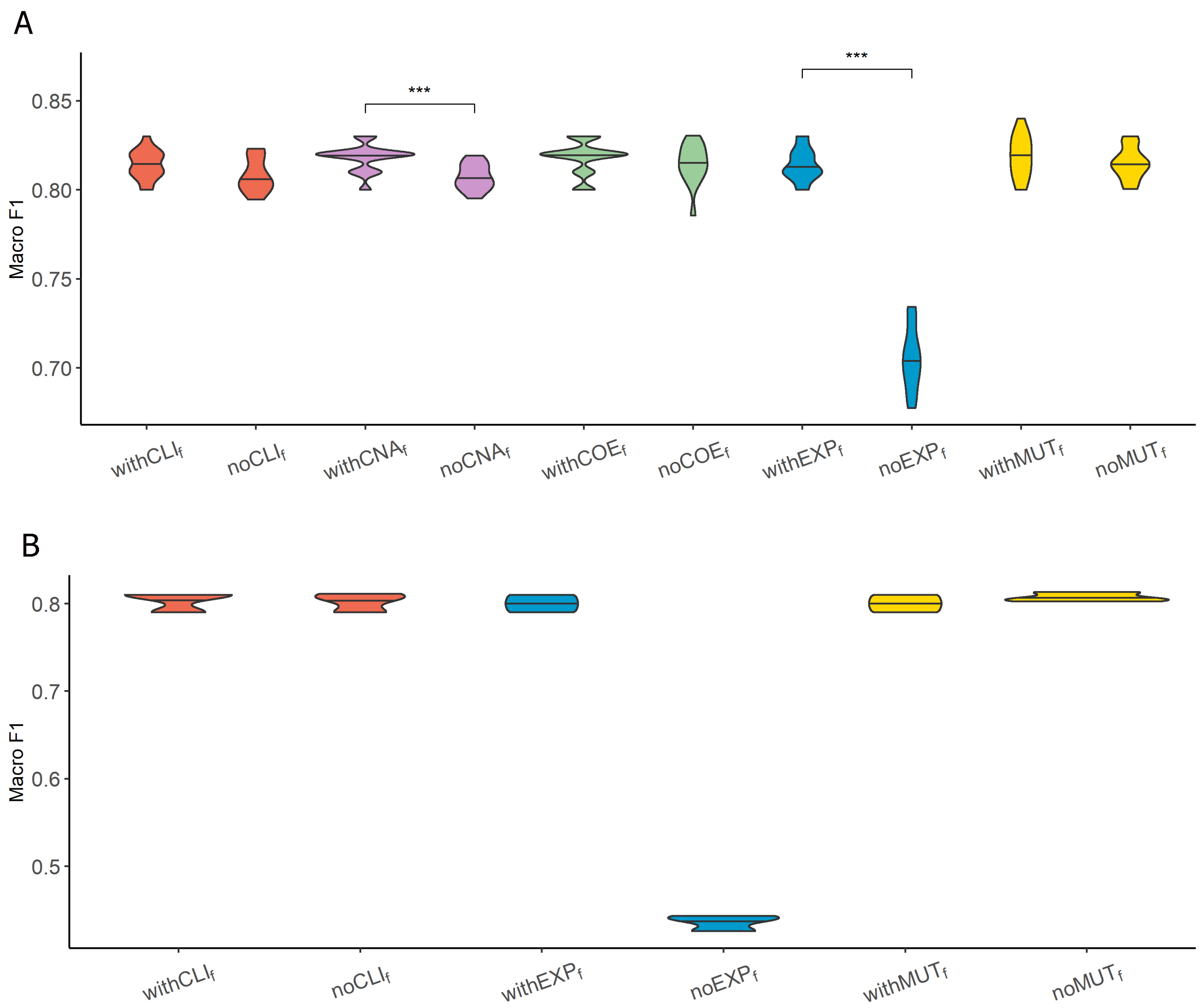


**Figure 8. Analysis of features from each datatype** Violin plot of macro F1 scores for the models excluding the features from each datatype ($no\mathbb{Y}_f$ models, where $\mathbb{Y}$ is the datatype whose features are completely excluded) versus corresponding SUPREME models ($with\mathbb{Y}_f$ models) on **(A)** METABRIC data and **(B)** the combined data. Significance level was measured between *with* and *no* cases of the same datatype (Wilcoxon rank-sum test p-value to compare the distribution of violin plots representing the significance $< 0.001$ by ***, else if $< 0.01$ by **, and else if $< 0.05$ by *). [Datatypes are abbreviated as CLI: clinical, CNA: copy number aberration, COE: coexpression, EXP: gene expression, MUT: mutation.]
